# Unifying population structure and relatedness analysis through a coalescent approach

**DOI:** 10.1101/2025.10.16.682899

**Authors:** Diego Veliz-Otani, Victor Borda, Heinner Guio, Omar Caceres, Cesar Sanchez, Carlos Padilla, Eimear E. Kenny, Sebastian Zollner, Timothy O’Connor

## Abstract

Standard methods in genome-wide association studies (GWAS) partition genetic similarity into recent familial relationship, modeled by a genetic relationship matrix (GRM), and distant relatedness, adjusted for using principal components (PCs). This practice relies on an implicit causal model that conflates population structure with confounding. Here, we challenge this approach by developing a unified framework grounded in coalescent theory. We introduce the Coefficient of Genealogical Similarity (GeSi), a statistic derived from a model of shared derived alleles that captures the full continuum of shared ancestry and can be estimated directly from genotype data. This leads to a new classification of GRMs into “full” matrices, which capture the complete genealogy, and “shallow” matrices, which measure only recent relatedness. Systematic benchmarking demonstrates that full GRMs are sufficient to model the genetic covariance from population structure, rendering PC adjustment for this purpose redundant. This finding clarifies that the justifiable role for PCs in such a model is to correct for true environmental or complex genetic confounders. Our analyses of empirical data confirm that including PCs can improve model fit, providing evidence that such confounding is present and correlated with axes of genetic variation. This work establishes a new theoretical framework that disentangles the modeling of genealogical relatedness from the correction of confounding, reframing the role of PCs as proxies for the latter and challenging the rationale for including the top PCs merely to capture maximal genetic variance.

## 1. Introduction

Genetic relatedness is a key concept in quantitative genetics, used to model phenotypic covariance and account for background genetic similarity in genome-wide association studies (GWAS) (1–3). Standard GWAS methods typically partition genetic similarity into recent, familial relatedness and distant, population ancestry (4,5). This approach is implemented by using a genetic relationship matrix (GRM) to model the genetic covariance due to recent relatedness and principal components (PCs) to regress out the effects of population structure (6,7). Here we argue that this partitioning is a methodological convenience, not a biological reality, and derive the coefficient of genealogical similarity (GeSi), which captures the full continuum of shared ancestry.

The biological effect of a given genetic variant on a complex trait is largely conserved across human populations. The average correlation of causal genetic effect sizes between individuals of European and African ancestry, for example, is r=0.98 across 53 quantitative traits in the UK Biobank (8). Of these 53 traits, 47 had a correlation that did not significantly differ from 1. Furthermore, the poor transferability of polygenic scores across populations can be largely explained by differences in linkage disequilibrium (LD) and allele frequency (9). This largely conserved biology of most quantitative traits contrasts with the standard practice of partitioning the genetic covariance into a component that should be modeled and one that should be regressed out.

The premise of discrete biological populations contradicts empirical data. Human genetic diversity is not organized into discrete clusters but is clinal and largely correlated with geographical distance (10–12). The perception of discrete clusters is often an artifact of biased sampling that over-represents geographic extremes and the misinterpretation of clustering software (12). Modern, diverse biobanks reveal a complex continuum of admixed ancestries that fills the gaps between previously defined reference populations, showing that participants exhibit gradients of genetic variation rather than distinct clusters (13).

This discrete view of human variation has led to a persistent conflation of population structure with confounding. From a theoretical standpoint, these concepts are different. Population structure is a source of genetic covariance arising from shared ancestry, which is a real biological signal that requires modeling (14). Classical (“environmental”) confounding arises from non-genetic factors, such as environment or culture, that are correlated with ancestry and also independently affect the phenotype (15). The term “genetic confounding” has been loosely used to refer to inflation of the test statistic due to distinct phenomena such as selection, assortative mating, and, population structure (16,17). However, the justification to consider these sources of statistical inflation (but not “recent” relatedness) as genetic confounding has not been justified.

The conflation of population structure with confounding is embedded in foundational methods. Genomic Control correctly identifies the causal mechanism of the inflation of the test statistic as genetic covariance due to population structure but refers to this inflation as confounding (14). LD-Score Regression explicitly groups population structure and cryptic relatedness into a single parameter for “confounding bias” and thus implicitly frames it as a source of inflation that should be corrected for (regressed out) rather than modeled (18). The distinction is critical, as standard corrections for genetic structure fail when the true confounder has a different spatio-temporal pattern than the genetic variation used for correction (19).

Collectively, this evidence questions the scientific basis for partitioning human genetic relatedness as the default *a priori*. Here we introduce the coefficient of Genealogical Similarity (GeSi) a statistic derived from coalescent model of shared derived alleles. We demonstrate that GeSi can be calculated directly from genotype data without inferring the genealogy. Extensive benchmarking of GeSi against other Genetic Relationship Matrices (GRMs) leads to a new classification of GRMs into “full” GRMs, which contain information of the full genealogy, and “shallow” GRMs, which measure recent relationships only. We show that under explicitly stated assumptions GeSi and other full GRMs are sufficient to model the full genetic covariance due to population structure without requiring principal components. This work establishes a new theoretical framework where environmental and genetic confounding are no longer conflated with population structure but are instead operationally defined as violations of specific assumptions, and where the role of principal components is re-framed as proxies for unmeasured confounders rather than as correction for population structure. While this work acknowledges the challenge of formally distinguishing these violations due to statistical identifiability, it provides a theoretical foundation upon which such methods can be developed.

## 2. Methods

### 2.1. Definition and generalization of the coefficient of genealogical similarity

We conceptualized the Coefficient of Genealogical Similarity (GeSi) from first principles, defining it first as a theoretical, genealogy-based metric (*GeSi*_*Tree*_) that represents the correlation in the number of derived alleles between two individuals based on their shared evolutionary history. We demonstrate that under an additive genetic model where causal variant effect sizes are independent of allele frequency (a model equivalent to the LDAK model with parameter *α* = 0), *GeSi*_*Tree*_ is also equivalent to the Pearson correlation of the individuals’ genetic values. Critically, we show that this coefficient can be accurately estimated from genotype data as the cosine similarity between raw genotype vectors, bypassing the need to infer the genealogy. We then use the estimator’s geometric interpretation to generalize GeSi by proposing that centering the genotype vectors on the ascertained sample’s mean and applying allele-frequency-dependent scaling factors (controlled by the LDAk’s *α* parameter) allows GeSi to model the correlation of genetic effects under more complex genetic architectures and relative to deviations from the ascertained sample mean rather than deviations from the ancestral phenotype. The full mathematical derivation and geometric extensions are detailed in the supplementary methods.

### 2.2. Data sources and preparation

We used both real and simulated data to extensively benchmark the performance of GeSi compared to standard GRM methods. All simulated phenotypes were quantitative. The real phenotype data corresponded to human height in the BioMe cohort. For each genotype dataset, we generated three variations: All biallelic single-nucleotide polymorphisms (SNPs) from whole-genome sequence (“WGS”), biallelic SNPs roughly in linkage-equilibrium (“LD-pruned”), and randomly sampled SNPs from the WGS data (“RS”), with the number of sampled SNPs matching that of the LD-pruned data. All datasets included autosomal variants only.

The datasets are fully described in the supplementary text. Briefly, the real datasets were a) The Peruvian Genome Project-Plus (PGP+, n=1,702), consisting of WGS data and demographic annotation; and b) WGS data from the BioMe cohort (n=9,885) and associated phenotype (height) and demographic metadata. We generated five sets of simulated data. Simulated set #1 (n=480) was an exploratory dataset consisting of WGS data generated using Msprime, and no phenotype data. Simulated sets #2-3 consisted of WGS data generated using Msprime and phenotype data simulated using R. Both simulated sets #2 and #3 consisted of four different scenarios, each of 5,000 samples. The four scenarios were: unrelated individuals from a homogeneous population resembling Iberians from Spain (IBS), unrelated individuals from an admixed population resembling the four Latino and Latin American (LAT) populations from the 1000 Genome Project (Colombians in Medellin, CLM; Mexicans in Los Angeles, MXL; Peruvians in Lima, PEL, and Puerto Ricans in Puerto Rico, PUR); related and unrelated individuals resembling IBS, and related and unrelated individuals resembling LAT. Simulated sets #4-5 (n=9,885) consisted of real WGS from BioMe and phenotypes simulated using R. The phenotypes were simulated *α* = 0 for Simulated Set #2, and *α* = -1 for all other simulated datasets (see supplementary methods). Simulated Set #5 included a confounder effect determined by the population labels; all other simulated datasets were generated without confounding effects. Simulated Set #1 consisted of one replicate and Sets #2-5 consisted of ten replicates.

### 2.3. Phenotype Simulation

We simulated quantitative phenotypes under a polygenic additive model under different conditions. The simulation strategies varied across three main axes, with full details provided in the supplementary methods. The relationship between minor allele frequency (MAF) and SNP effect size was modeled using the LDAK framework’s *α* parameter, focusing on the *GeSi*_*tree*_ model (*α* = 0) and the standard GCTA model, which is a special case of the LDAK model with *α* = -1. For Simulated Set #5, we introduced an environmental confounding effect determined by the population labels in the BioMe cohort (AFR, EUR and LAT).

The simulated traits had a heritability of 0.5 determined by 1000 causal sites. We explored other values of heritability (0.2 and 0.8) and polygenicity (100 and 10,000 causal sites) in the initial simulations; however, the results were largely the same and dropped them from further analyses to save computational time (results not shown). Causal variants were selected with uniform probability along the genome from WGS data, and their effect sizes were simulated from random normal distribution with mean 0 and variance 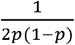, where *p* is the MAF of the causal site. Finally, we added a residual variance so that the empirical heritability matched the target heritability of 0.5 (see supplementary methods).

### 2.4. Calculation of the genetic relationship matrices

The procedure to calculate GeSi and the standard genetic relationship matrices (GRMs) GCTA/LDAK and PC-Relate is fully described in the supplementary methods. Briefly, the GeSi and LDAK matrices were computed using all three genotype data variations: WGS, LD-pruned, and randomly sampled (RS) sites, with *α* values ranging from -2 to 0 in increments of 0.25. The LDAK matrix with *α* = -1 is equivalent to the GCTA GRM. Due to its high computational requirements, the PC-Relate matrix was calculated only with *α* = -1 and only from LD-pruned data, as is standard for the method, using the KING-robust GRM as an initial step in its iterative calculation pipeline.

### 2.5. Mixed Linear Model association test

We performed all association tests and variance component estimations using mixed linear models (MLMs) as implemented in the GENESIS package for R. The general inference model was Y= Xβ + Zg + ∈, where the covariance of the random genetic effects vector *g* was defined either by a GeSi, LDAK/GCTA or PC-Relate GRM matrix. To assess the impact of ancestry correction, all models were run with either zero or ten principal components (PCs) as fixed-effect covariates. The MLMs for the Msprime-simulated data (Simulated Sets #2 and #3) only included PCs as covariates. The MLMs for the BioMe-based simulations (Sets #4 and #5) and the real BioMe phenotype data were further adjusted for sex, age, age^2^ and demographic label.

We assumed homoscedastic residuals for simulated data analyses without confounding (Simulated Sets #2-4). For the simulation set that included confounding effects (Set #5), we allowed for heteroscedasticity by fitting a different residual variance component for each demographic label. For the real BioMe height data, a residual variance component was fit to each of the sex-by-population groups.

### 2.6. CV-BLUP-based predictions

We used the Best Linear Unbiased Predictor (BLUP) equation(1,20) to predict the total genetic effects (also known as the breeding value in plant and animal genetics) in the held-out sets of a 20-fold cross-validation (CV) analysis of Simulated Sets #4 and #5, and the actual height values from the BioMe cohort. Because the true *α* value is unknown in real data, we repeated this analysis using *α* values from -2 to 0 in increments of 0.25. This allowed us to both find the *α* value that maximizes the CV-BLUP prediction accuracy in the BioMe cohort, and to assess the sensitivity to *α* value misspecification in the simulated datasets.

All datasets were split into 20 cross-validation folds of 494 subjects each. Because the total sample size (9,885) was not a multiple of 20, we randomly excluded the same five samples from all cross-validation analyses. We used the `GENESIS::fitNullModel` function to fit a mixed linear model on the training set of each cross-validation fold. The null models included a genetic similarity matrix (either a GRM or a GeSi matrix) and the same covariates as in the MLM association testing. We obtained the variance components estimates for each fold, which were then fed into the BLUP equation:

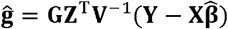

Where **Y** ∈ ℝ^*n*×1^ is the vector of phenotypes of the training set (n = 9,386); **X** ∈ ℝ^*n*×*p*^ is the matrix of fixed effects covariates in the training set 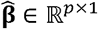 is the vector of fixed effects coefficients estimated in the training set; 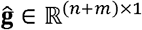 is the predicted genetic effects of both the training and validation (m = 494) sets; 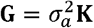 is the estimated covariance matrix of the genetic effects; **K** is the genetic similarity matrix of both training and validation sets, and **G, K** ∈ ℝ^(*n*+*m*)×(*n*+*m*)^. The incidence matrix **Z** ∈ ℝ^*n*×(*n*+*m*)^ maps the genetic effects to each subject. **V**= (**ZGZ**^T^ + **R**), where **V** ∈ ℝ^*n*×*n*^ is the total variance matrix, and **R** ∈ ℝ^*n*×*n*^ is the diagonal matrix of residual variance.

We calculated the total phenotype of the validation set as the sum of estimated fixed effects and random genetic effects 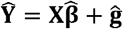. For each CV fold, we measured the accuracy of the genetic value and phenotype prediction in the held-out data via the squared Pearson correlation coefficient (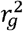 and 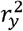, respectively) and the coefficient of determination defined as 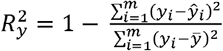, and analogously, 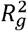. For each data set, GRM type algorithm (LDAK/GCTA or GeSi), and data type (WGS, LD-pruned or RS), we found the *α* value that minimized the 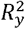 in the real BioMe data.

## 3. Results

We start by first validating the genealogical interpretation of the original *GeSi*_*tree*_ definition. We then validate the two generalizations of the *GeSi*_*tree*_ model, namely that the frame of reference can be set to that of mean phenotype in the ascertained samples by centering the genotype matrix in the sample mean, and that by changing the scaling factor of GeSi’s denominator, we can allow for an allele frequency-dependent distribution of effect sizes, similar to that of LDAK’s approach. We demonstrate the validity of these extensions by extensively benchmarking GeSi against LDAK/GCTA and PC-Relate under different conditions in Genome Wide Association Studies (GWAS), heritability estimation. We then test GeSi’s and other GRMs’ ability to predict phenotypes in a BLUP framework to assess whether including PCs in the mixed linear model adds any information that would be translated into an increase in predictive accuracy. Finally, we use GeSi and other GRMs to estimate the heritability of height in the BioMe data.

### 3.1. Definition of GeSi and exploration of its basic properties

We derived an expression that measures the pairwise correlation of number of derived alleles in terms of the evolutionary times. We call this expression GeSi, short for coefficient of Genealogical Similarity. We then demonstrate that under three specific assumptions, GeSi also represents the correlation of phenotypic deviations from the phenotype of an ancestor used to define the ancestral allele states. The general framework is illustrated in **Fig. 1a-e**, and the formal definitions and derivations are detailed in the **Methods** and **Supplementary text**. The three assumptions are: 1) genetic effects are purely additive; 2) genetic effects accumulate linearly with time; 3) genetic effects are independent of environmental effects. This framework leads to the following operational definitions: 1) “genetic confounding” as the violation of the second assumption due to, for instance, assortative mating or selection, but not due to population structure, whose effect on the genetic covariance is modeled by GeSi; and 2) “environmental confounding” as a violation of the third assumption, either by a direct correlation of environmental effects with the polygenic score, or by an indirect correlation with the polygenic score via correlation with the ancestry components.

**Figure 1.**
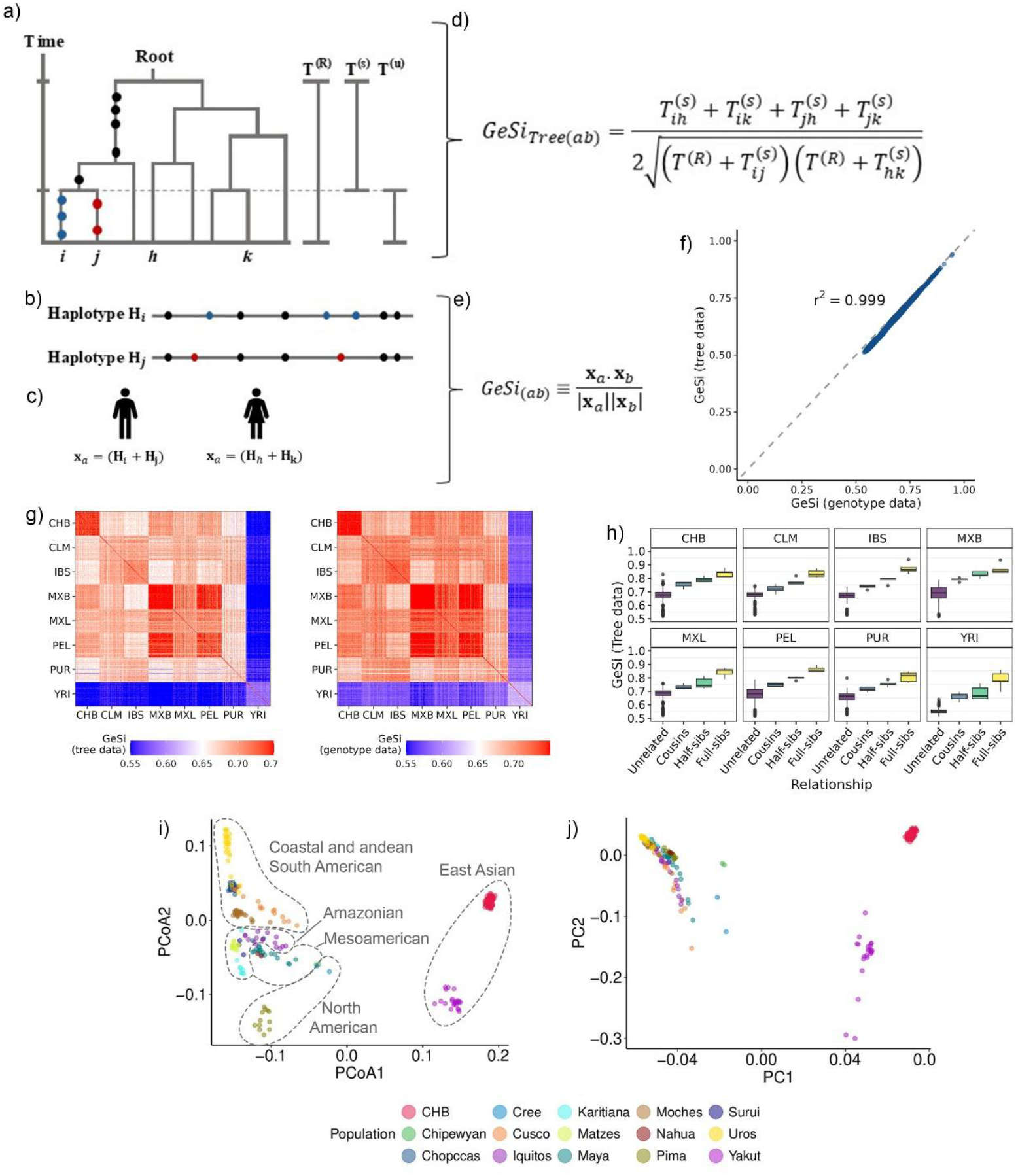
Parameterization and general features of GeSi. **a)** Genealogy of four arbitrary haplotypes *i, j, h* and *k*. The circles on the tree represent the mutations carried by haplotypes *i* and *j* as they occurred across time; mutations on other haplotypes are not shown. ***T***^(***s***)^: Shared divergence time between haplotypes *i* and *j*. ***T***^(***u***)^: Unshared divergence time between haplotypes *i* and j. ***T***^(***R***)^: Time from the present to the root of the tree. **b)** Schematic showing the distribution of the mutations across either or both haplotypes *i, j* depending on when they occurred. The mutations marked with a black circle are shared by both haplotypes. The mutations marked with a blue or red circle occurred after their genealogies diverged and are exclusively found on either haplotype. **c)** Genotype vectors **x** expressed as the sum of the haplotype vectors **H.** The genotype vectors of two subjects carrying the pair of haplotypes (*i, j*) and (*h, k*) are indexed by the subscripts *a* and *b*, respectively. Thus, the genotype vectors are the sum of the haplotype vectors: **x**_***a***_ = (**H**_***i***_ **+ H**_***j***_) and **x**_***b***_ = (**H**_***h***_ **+ H**_***k***_). **d)** and **e)** connection between the time parameters and the genotype vectors and the GeSi coefficient and its estimator, respectively. **f)** Comparison of the tree-based and genotype-based GeSi estimates shows a high correlation between both values. **g)** Heatmaps of the GeSi values, clustered by populations labels, calculated from the true genealogical trees (left heatmap) and from genotype data (right heatmap) in Simulation set #1 (n=480, see methods). Both estimates reveal similar patterns of population structure corresponding with the simulated population labels. **h)** Boxplot of the tree-based GeSi values of same-population pairs stratified by familial relationship and population of origin. **i)** Principal Coordinate Analysis (PCoA) of the genotype-based GeSi values transformed into angles via an arc-cosine transformation. **j)** First two principal components estimated via Plink. r^2^: Squared Pearson’s correlation coefficient.

We proved that the cosine-similarity of biallelic genotype vectors is an estimator of GeSi that can be calculated directly from biallelic genotype data without knowledge of the genealogical trees (**Fig. 1b, c, e, f**). Next, we show that both the tree-based definition of GeSi and its genotype-based estimates reveal similar patterns of population structure in the Simulation Set #1 (**Fig. 1g**). The results of **Fig. 1f** and **1g** confirm the suitability of using the genotype-based estimator for all remaining analyses. Next, we visualized the distribution of *GeSi*_*tree*_ values stratified by population and familial relationship level (**Fig. 1h**) and confirmed that GeSi is not only indicative of population structure, but also of recent relatedness. Furthermore, the distribution of GeSi values between intra-population pairs of unrelated subjects differed across populations (**Fig. 1h**), highlighting that the basal genetic similarity between “unrelated” individuals is different across populations, in concordance with the definition of *GeSi*_*tree*_, which is inversely proportional to the genome-wide average coalescence time.

Since the genotype estimator of GeSi represents the cosine similarity of the genotype vectors, we considered that taking its arc-cosine would yield a natural measure of distance. Thus, we took the arc-cosine of the GeSi matrix and show that its PCoA projection better resolves the structure of Pan-American populations than a conventional PCA (**Fig. 1i** and **1c**).

### 3.2. Fully informative GRMs do not require principal components to adjust for admixture in single-variant association tests in the absence of confounding

The purpose of this simulation was two-fold, first, to validate the generalizations of GeSi by mean-centering the genotype matrix and changing the scaling factor *α* to values other than zero, and second, to assess whether PCs are required to adjust for inflation due to population structure or admixture. We benchmarked the generalized GeSi estimators against standard methods in single-variant association tests using the Msprime-generated Simulation Sets #2 (*α* = 0) and #3 (*α =* -1).

The GeSi and GCTA/LDAK GRMs accurately estimated SNP effect sizes across all scenarios, with slightly higher accuracy in admixed (LAT) samples than in homogeneous (IBS) ones. Including principal components (PCs) as covariates in models with GRMs built from dense data WGS or random sites (RS) consistently reduced the accuracy of effect size estimates by introducing a downward bias (slope falling farther away from 1) and reducing the Pearson’s correlation between true and estimated effect sizes. This effect was absent for models using GRMs built from LD-pruned data (**Suppl Fig. 1 and 2**). The PC-Relate GRM, built from LD-pruned data, yielded effect size estimates with accuracy comparable to that of other LD-pruned GRMs and was also unaffected by the inclusion of PCs (**Suppl Fig. 1 and 2**). Detailed results are shown in **Suppl. Tables 2-3**.

Test-statistic inflation was primarily influenced by the data type used to construct the GRM. Models using WGS-based GRMs were the most conservative (mean *λ*_*gc*_ ≤ 1.0), whereas those using LD-pruned GRMs showed slight inflation (mean *λ*_*gc*_ ≈ 1.03 − 1.07) **Suppl. Fig. 3 and 4**.

These simulations lacked non-genetic confounding, and the inflation was observed in all four demographic and sampling scenarios. Thus, the observed inflation does not reflect uncorrected population or confounding, but rather the expected polygenic signal from non-causal variants tagging true causal variants. While PC inclusion did not affect inflation for GeSi or GCTA/LDAK models, it was essential for controlling inflation in PC-Relate models, particularly in the admixed and relatedness scenarios, where models without PCs showed significantly greater inflation (**Suppl. Fig. 3 and 4**).

Finally, a trade-off between inflation control and statistical power was observed. Models with LD-pruned GRMs showed slightly higher raw power to detect causal variants than models using WGS or RS-based GRMs (**Suppl. Fig. 5 and 6**). However, after adjusting for test-statistic inflation via the genomic control, the statistical power was equivalent across all methods and modeling choices.

### 3.3. Heritability estimation requires information-dense GRMs but not principal components in the absence of confounding

Across both the *α* = 0 (Simulation Set #2) and *α* = -1 (Simulation Set #3) scenarios, heritability estimation was highly sensitive to the information density of the GRM but mostly unaffected by the inclusion of PCs as covariates. GRMs calculated from WGS data consistently produced the most accurate and least biased heritability estimates. In contrast, GRMs built from either LD-pruned or randomly sampled (RS) sites resulted in significant downward bias (**Figure 2**, and **Suppl. Fig. 7-9**). Detailed numerical values are shown in **Suppl. tables 4 and 5** for Simulation Sets #2 and #3, respectively. The *α* = -1 simulation allowed for a direct comparison with PC-Relate, which yielded the most downward-biased heritability estimates of any method tested. Even when compared to GeSi and GCTA/LDAK matrices built from the same LD-pruned data, PC-Relate’s estimates were significantly lower, and including PCs as covariates did not correct this severe downward bias (**Fig. 2**, and **Suppl. Fig. 7-9**). This suggests that PC-Relate, by design, fails to capture the heritability component attributable to distant relatedness, which is removed when regressing the genotypes on the top principal components.

**Figure 2.**
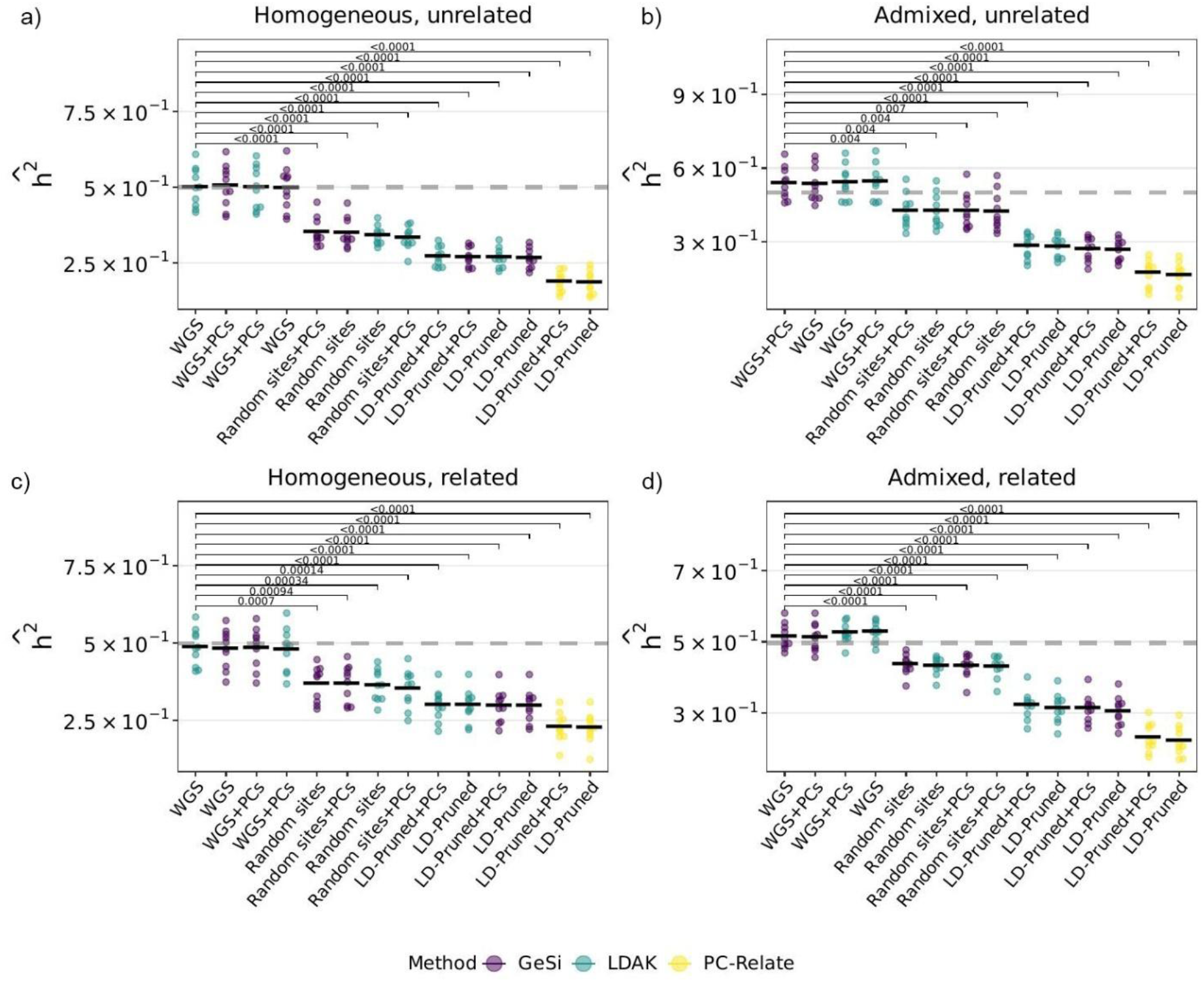
Squared error of the heritability estimates under different levels of recent relatedness and admixture when the true scaling factor is *α* = −1 in the Simulation Set #3. The X axis details whether the mixed linear model (MLM) used a GeSi, LDAK or PC-Relate genetic relationship matrix, and whether principal components (PCs) were included in the model. All GRMs were calculated with *α* = −1. Ten replicates are shown for each model. Each GRM was calculated from three different types of color-coded genetic data: Whole Genome Sequence (WGS) data, LD-pruned data, or randomly selected sites. Each panel represents one of the four different combinations of demographic and ascertainment scenarios that were simulated: **a)** Unrelated individuals from Iberians in Spain (IBS); **b)** Mixture of related and unrelated individuals from IBS; **c)** Unrelated individuals from Latino and Latin American populations (LAT; see main text and methods); and **d)** Mixture of unrelated and unrelated individuals from LAT. A two-sided Wilcoxon rank sum test was used to compare all methods against the MLM model with 10 PCs and an LDAK matrix calculated from LD-pruned data. Models are displayed from best to worst within each panel as judged based on the mean squared error. A square bracket and FDR-adjusted p-value are shown for all comparisons with an FDR <0.05. Black horizontal lines: Mean squared error.

### 3.4. Heritability estimation in the presence of simulated confounding

To validate our findings on more realistic genetic data and to assess the impact of confounding, we simulated phenotypes using genotypes from the BioMe cohort, first without (Simulation Set #4) and then with (Simulation Set #5) an environmental confounder correlated with population labels

The analysis of the non-confounded baseline data (Set #4) largely replicated the patterns observed in the Msprime simulations. Heritability estimates were most accurate when GRMs were calculated from WGS data, while using LD-pruned or RS data led to significant underestimation **(Suppl. Fig. 10a** and **10b**). A notable difference emerged in the direction of bias for WGS-based GRMs: WGS-GCTA produced moderately upward-biased estimates, whereas WGS-GeSi was slightly downward-biased. As before, PC-Relate yielded the most severely underestimated heritability, and including PCs as covariates did not improve the accuracy for any GRM type in this non-confounded scenario (**Suppl. Fig. 10a** and **10b**).

These patterns remained consistent after introducing the environmental confounder (Set #5). The relative performance of the methods and data types was unchanged, with WGS-GRMs remaining superior and PC-Relate performing the worst (**Suppl. Fig. 10c** and **10d**). Crucially, even in the presence of a confounder determined by the demographic labels, the heritability estimates of the models with either a GeSi or GCTA GRM were not affected by the inclusion of PCs.

### 3.5. Phenotype prediction using fully informative and partitioned GRMs in a BLUP-framework

To directly test the information content of each GRM, we used BLUP to predict both the total genetic effects (*g*) and the phenotype (*y*) in the BioMe-based Simulation Sets #4 and #5. In the absence of confounding (Simulation Set #4), GeSi and GCTA GRMs were the most accurate predictors of the true genetic value, particularly when constructed from WGS data. In contrast, PC-Relate was by far the least accurate predictor of the total genetic effects *g*, confirming that it contains less genealogical information (**Fig. 3a**). For predicting the total phenotype (Y), however, the performance of PC-Relate improved to the level of other LD-pruned GRMs, but only when PCs were included as covariates (**Fig. 3b**). Conversely, phenotype prediction accuracy for GeSi and GCTA models was not improved by the addition of PCs (**Fig. 3b**). In fact, the accuracy of the GCTA models calculated from WGS data or random sites dropped after adding PCs to the model. Similarly, all MLMs using either a GeSi or GCTA GRM lost accuracy at predicting the genetic effects after adding PCs to the MLM (**Fig. 3a**). This demonstrates that the information contained in the top PCs is essential for MLMs using a PC-Relate GRM but redundant if the MLM uses a GeSi or GCTA/LDAK.

**Figure 3.**
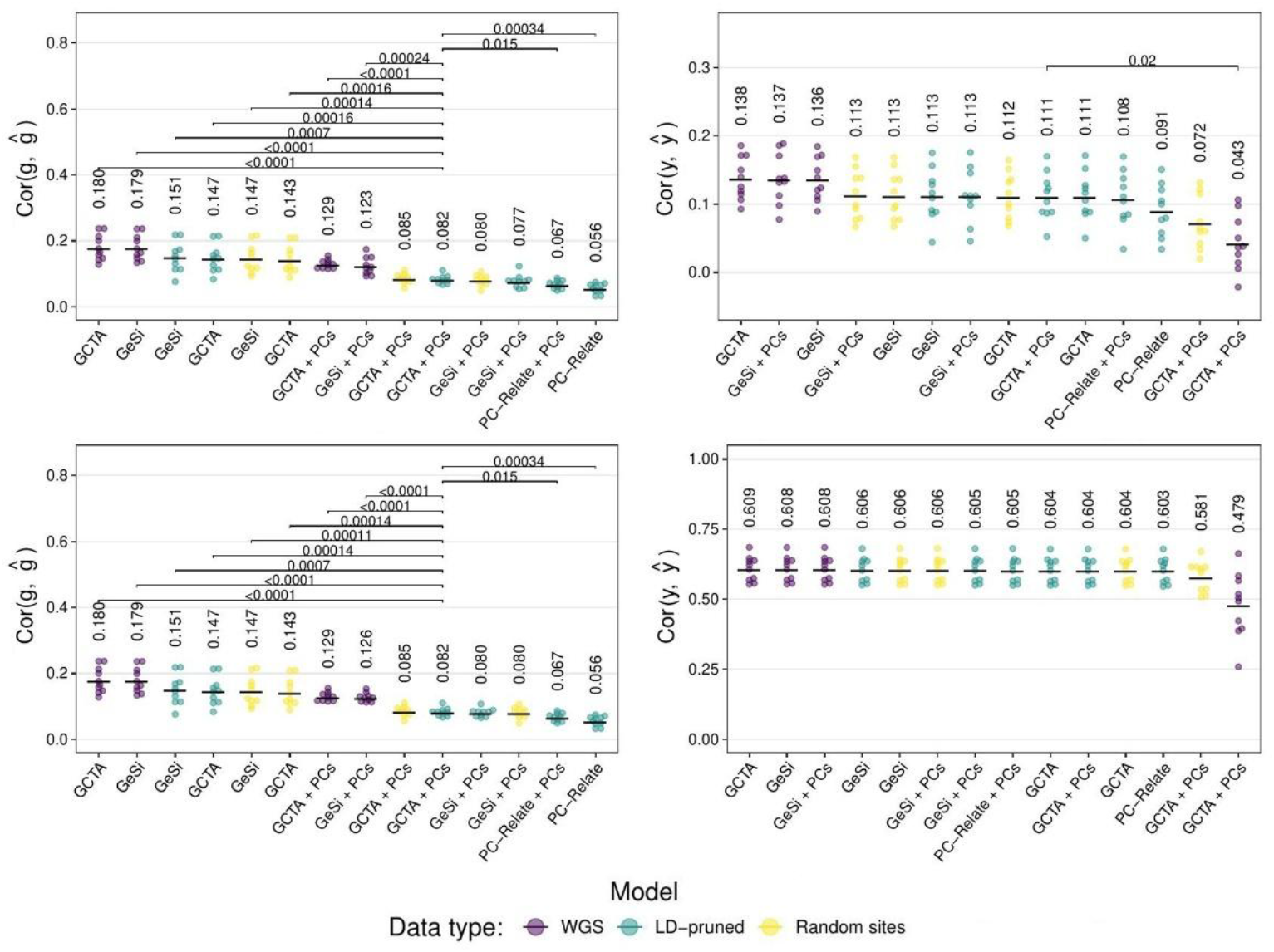
Accuracy of the predicted genetic value 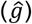 and phenotype (*ŷ*) via BLUP using different mixed linear model specifications in Simulated Set #4. Panels **a** and **b** correspond to Simulation Set #4 (no confounding), and panels **c** and **d**, to Simulation Set #5 (confounding determined by population labels). Panels **a** and **c** show the accuracy of the genetic value (g) prediction, and panels **b** and **d** the accuracy of the phenotype prediction. Each point represents the result of a cross-validation fold. The horizontal axis details whether the mixed linear model (MLM) used a GeSi, LDAK or PC-Relate genetic relationship matrix (GRM), and whether principal components (PCs) were included in the model. All GRMs were calculated with *α* = −1. Each GRM was calculated from three different types of color-coded genetic data: Whole Genome Sequence (WGS) data, LD-pruned data, or randomly selected sites. A two-sided Wilcoxon rank sum test was used to compare all models against an MLM with 10 PCs and a GCTA matrix calculated from LD-pruned data. Models are displayed from best to worst within each panel as judged based on the mean squared error. A square bracket and FDR-adjusted p-value are shown for all comparisons with an FDR <0.05. Black horizontal lines: Mean-squared error

These findings remained virtually identical in the presence of a known environmental confounder that was included in the model (Simulation Set #5, **Fig. 3c** and **Fig. 3d**). The predictive accuracy for the genetic value u followed the same pattern, with GeSi and GCTA GRMs outperforming PC-Relate (**Fig. 3c**). In this simulation, however, the strong effect of the confounder itself (which was included as a covariate in all models) drove a large portion of the phenotypic prediction accuracy (**Fig. 3d**), which had the effect of largely masking the performance differences between models that were apparent in the non-confounded scenario, and in the genetic value prediction of the confounded scenario (**Fig. 3c**).

### 3.6. Power to detect population structure via principal components

A potential critique of our finding that PCs are redundant when used with holistic GRMs is that the sample size may be insufficient for the PCs to capture population structure. To formally address this, we evaluated whether our sample provides adequate statistical power based on the spectral theory of Patterson et al (21) and Bryc et al (22). Although defining populations in our admixed cohort is artificial, we can directly test our power to detect the specific structure that was explicitly simulated as the source of the confounder in Simulation Set #5.

We first quantified the differentiation between the population labels (AFR, EUR, LAT) in the unrelated subjects of the BioMe cohort (n = 9,190), which yielded an F_ST_ of 0.024. Using the calculated F_ST_ to parameterize the detection threshold as (1 + *F*_*ST*_)/2, as proposed by Bryc et al., we obtained a significance threshold of *t* = 0.512 for the normalized eigenvalues of the uncentered genotype covariance matrix. Our analysis of the BioMe genotypes revealed seven normalized eigenvalues above this threshold (1986.7, 47.8, 8.1, 2.0, 1.6, 1.1, and 0.8).

This result demonstrates that our sample size provides more than sufficient statistical power to detect the genetic structure associated with the labels used to generate confounding in our simulations. Therefore, the observed redundancy of PCs in the preceding heritability and BLUP analyses is not an artifact of low power. This strengthens our conclusion that full GRMs, such as GeSi and GCTA, already contain the structural information captured by the top PCs. This is further corroborated by the consistently poor performance of the structure-corrected GRM from PC-Relate when PCs are not included in the mixed linear models.

### 3.7. Prediction of human height in real data from the BioMe cohort

To test whether PCs would also be unnecessary to predict phenotypes when there are unknown sources of confounding, we predicted human height in the BioMe cohort. Unlike our simulations where confounders were known, here we included demographic labels as fixed-effect covariates, which serve as imperfect proxies for the true, unobserved environmental and social factors that may be partially correlated with ancestry. Because the true total genetic effects are unknown, we could only measure the accuracy of the phenotype prediction, which was assessed via the coefficient of determination (R^2^) in a 20-fold cross-validation analysis. Because the true underlying genetic architecture (i.e., the parameter *α*) is unknown for real traits, we first compared all methods using a fixed value of *α* = -1 before identifying the optimal value for each method.

When *α* was fixed at -1, the results resembled those observed in our simulations without confounding. Specifically, most models using GeSi or GCTA GRMs performed similarly well, with the notable exception of the GCTA-WGS model, which lost accuracy when PCs were included in the model (**Suppl. Fig. 11a**). The PC-Relate model without PCs had the lowest predictive accuracy of all tested methods. However, its accuracy improved substantially, becoming comparable to other LD-pruned GRMs, after including PCs as covariates (**Suppl. Fig. 11a**).

These patterns were largely replicated when selecting the alpha value that maximized the cross-validation predictive accuracy for each method, although PC-Relate was kept fixed at *α* = -1. (**Suppl. Fig. 11b**). In this optimal scenario, all models using either a GeSi or LDAK GRM achieved similar predictive accuracy, with a slight, consistent but non-significant trend of WGS models to achieve higher accuracy than other models. For these methods, the inclusion of PCs as covariates had no impact on the predictive power. PC-Relate remained the only method that demonstrated a clear benefit from PC adjustment, and its model without PCs was the poorest performer overall. This confirms that even in a real-world setting with complex and unknown confounding, the information captured by the top PCs is redundant for GRMs that do not partition the genetic relatedness, but essential for partitioned GRMs like PC-Relate. Full results are presented in **Suppl. Table 6**.

### 3.8. Heritability estimation of human height in the BioMe cohort

We next estimated the heritability (h^2^) of height in the BioMe cohort to test our conclusions on a real complex trait where the true genetic architecture and sources of confounding are unknown. Because the optimal alpha value is not known for real traits, and because its misspecification can impact the accuracy of h^2^ estimates (**Suppl. Fig. 12**.), we performed an alpha scan for GeSi and LDAK models. We selected the alpha for each method that minimized the Akaike Information Criterion (AIC) of the MLM, which included sex, age, age^2^, population label, and sex-by-population label interaction terms as covariates, and accounted for variance heteroscedasticity across sex-by-population groups. We ran all models in duplicate, with and without ten principal components as covariates. The GRMs were calculated with alpha values ranging between -2 and zero in increments of -0.25 (**Suppl. Fig. 13**).

The model with the best overall fit used a GeSi matrix calculated from WGS data with an optimal alpha of -0.5 and included PCs as covariates. Because we allowed for heterogeneous residual variance among the sex-by-population groups, we obtained a different heritability estimate for each of them. **Figure 4** shows the sample size-weighted average heritability across all six groups sorted left to right from lowest to highest AIC. The group-specific estimates are presented in **Suppl. Fig. 14**.

**Figure 4.**
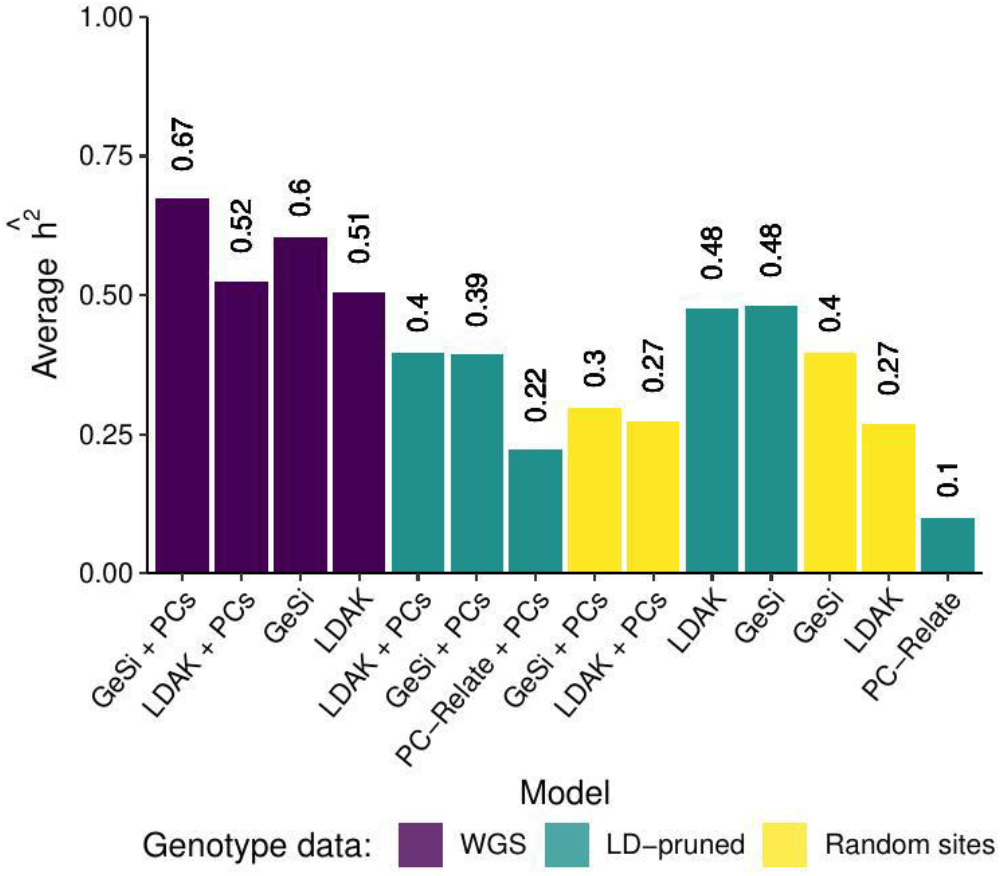
Average heritability estimates of human height. We identified the value of the *α* parameter that minimized the AIC of each model defined on the horizontal axis and data type indicated in the legend. The models are sorted from lowest to highest AIC from left to right. We allowed for heterogeneous residual variance across each of the sex-by-population groups, and thus obtained six different heritability estimates for each model for each data type. The vertical axis shows the sample size-weighted average heritability estimate of the six sex-by-population groups. The group-specific estimates are shown in **Suppl. Fig. 14** (estimates with the *α* value that minimizes the AIC) and **Suppl. Fig. 15** (*α* fixed at -1). The selection of the *α* parameter value is detailed in **Suppl. Fig. 13**.

Consistent with our simulations, models using GRMs calculated from WGS data yielded higher h^2^ estimates than GRMs calculated from either LD-pruned or RS data. Models that included PCs consistently achieved a better fit than those without them (**Suppl. Fig. 13**). This suggests the PCs captured variance from unmeasured confounders not fully accounted for by the available demographic labels. For a direct comparison with PC-Relate, we also estimated heritability with *α* fixed at -1 (**Suppl. Fig. 15**). In this analysis, the results mirrored our simulation findings, namely, that after even after controlling for type of genotype data, PC-Relate produced the lowest heritability estimates.

However, we also considered the possibility that the PCs did not capture unknown confounders, but instead simply captured the axes of greatest variation more efficiently than the GRMs. To assess this hypothesis, we defined Δ*AIC* = *c* where negative Δ*AIC* values mean that the model improved by adding PCs. We calculated the average Δ*AIC* across the ten replicates in Simulation Set #4 (without confounding, **Suppl. Table 7**), Simulation Set #5 (with a known confounding, **Suppl. Table 8**), and the Δ*AIC* on the real height data from BioMe (**Suppl. Table 9**). In neither simulation did adding PCs improve the model fit (average Δ*AIC* > 0) for GWAS models with a GeSi or LDAK/GCTA matrix calculated with the true or close to the true *α* parameter value (-1.5 < *α* ≤ 0). In contrast, and in accordance with **Suppl. Fig. 13**, adding PCs to GWAS models of the true height data resulted in negative Δ*AIC* values, indicating that adding PCs improved the model fit. Finally, adding PCs to GWAS models with a PC-Relate matrix improved the model fit in both Simulation Sets and in the real height data. These results suggest that the improvement of the model fit in the real height data by adding PCs to GWAS models with a full GRM is caused by the PCs’ role at correcting for either genetic or environmental confounding rather than by more efficiently capturing genetic covariance due to population structure than the full GRMs.

## 4. Discussion

Herein we introduced GeSi, a new metric derived from a coalescent theory point of view that models genetic relatedness as a continuous measure of shared genealogical history. We first demonstrated that the genotype-based cosine similarity is an accurate estimator of the theoretical, tree-based definition of GeSi (**Fig. 1f**), successfully capturing both deep ancestral relationships and recent familial kinship in a single matrix (**Fig. 1f-i**). By providing a continuous measure of genealogical similarity, GeSi directly addresses the limitations of conventional IBD-based metrics, which disregard the variable background similarity among individuals classified as “unrelated” (**Fig. 1h**).

Our subsequent benchmarks reveal that genetic relationship matrices can be broadly categorized based on the information they contain. Full GRMs, such as GeSi and the standard GCTA and LDAK matrices, retain information of the full genealogical history. In contrast, shallow GRMs, like PC-Relate (4), are designed to discard ancestral relatedness information and therefore contain less genealogical information than full GRMs. We selected PC-Relate as a representative of shallow GRMs because, it is an accurate estimator of the kinship coefficient even in highly structured and admixed populations (23). Other methods for estimating recent kinship, such as those based on inferred identical-by-descent (IBD) segments, have been shown to have an accuracy similar to or higher than that of PC-Relate at estimating the kinship coefficient (23). Although we did not include IBD-based GRMs in our benchmarks due to their computational cost, it is reasonable to infer that any estimator focusing strictly on recent relatedness would be less information-rich than a full GRM.

Our results consistently show the downstream effects of this distinction between full GRMs and shallow GRMs. The heritability was severely underestimated by PC-Relate, which fails to capture the variance component attributable to distant relatedness (**Fig. 2**). Furthermore, our BLUP analyses showed that full GRMs consistently outperformed PC-Relate at predicting an individual’s total genetic value (**Fig. 3a** and **3c**). Moreover, neither the phenotype prediction (**Fig. 3b** and **3d**) nor the total genetic value prediction (**Fig. 3a** and **3c**) improved after adding principal components to mixed linear models containing a full GRM. Therefore, full GRMs do not require principal components as covariates to account for population structure in mixed linear models, whereas the performance of PC-Relate, and presumably other shallow GRMs, is fundamentally dependent on the inclusion of PCs (**Fig. 3**). The notion that a single GRM can model both recent and distant relatedness is further supported by recent evidence showing that genetic effect sizes for most complex traits are highly conserved across continental ancestries (8).

A potential limitation of our findings is that the redundancy of principal components when used with a full GRM could arise from insufficient statistical power to detect population structure. We formally evaluated this possibility using the spectral theory of eigenanalysis (21,22). Although these frameworks are built on simplified models of discrete, non-hierarchical populations, we can apply them to test our power to detect the specific structure that was explicitly simulated as a confounder in our analyses. Using the provided continental labels to define populations, we calculated a between-group divergence of *F*_*ST*_ = 0.024. According to the theory proposed by Bryc et al., this corresponds to a significance threshold of *t* = 0.512 for the normalized eigenvalues of the uncentered genotype covariance matrix. Our analysis revealed seven eigenvalues (1986.7, 47.8, 8.1, 2.0, 1.6, 1.1, and 0.8) that exceeded this threshold, confirming that our sample provides sufficient power to resolve the major axes of genetic variation. Importantly, the entries of the GCTA GRM are functions of the expected pairwise coalescence times (24), and the PCs of the genotype matrix correspond to the eigenvectors of the GRM (24,25), which further supports the notion that both the full set of PCs and the GCTA GRM contain the same information about the full genealogy.

Our new framework of studying genetic relatedness as a continuum leads to a more nuanced interpretation of the role of PCs in practice. The common justification for including PCs is to correct for confounding, but this rests on the central paradox that the application of PCs often inverts the nature of the effects being modeled. Theoretical frameworks that give genetic meaning to PCs rely on unrealistic models of discrete populations (22,26), yet the complex, non-linear effects of the social environment are then modeled as a simple linear functions of those PCs. This paradox arises from an informal but pervasive conflation of population structure with confounding itself (15). From a coalescent perspective, population structure is simply a source of genetic covariance that a full GRM can appropriately model under certain conditions. The true confounding that biases association studies occurs when non-genetic factors, such as social constructs, cultural practices, and environmental exposures, are correlated with the genetic ancestry that PCs measure (27,28). Furthermore, adjusting for PCs can introduce statistical artifacts like collider bias, if the PCs capture local genomic features rather than genome-wide ancestry (29). Despite these limitations, we acknowledge that PC adjustment has been, in practice, an effective strategy for mitigating inflation due to confounding.

The improved model fit from including PCs in the real-data analysis of height initially raised a question of statistical identifiability. The improvement could stem from PCs acting as proxies for unmeasured confounders, or from PCs simply capturing the axes of greatest genetic variance more efficiently than a full GRM. Our Δ*AIC* analysis provides a direct test of these alternatives (**Suppl. Tables 7-9**). In simulations without confounding (Simulation Set #4) or with a known confounder (Simulation Set #5), adding PCs to a model with a full GRM consistently worsened the model fit (Δ*AIC* > 0). This finding provides strong evidence against the hypothesis of statistical efficiency. It supports the conclusion that the utility of PCs in the real-data model stems from their role in capturing variance from true environmental (19) or genetic confounding that were present in the cohort but absent from our baseline simulations.

This evidence sharpens the critique of current practices for PC selection. Current practice often involves the arbitrary inclusion of a fixed number of top PCs (e.g., 10 or 20), an atheoretical approach with no guarantee of capturing the relevant confounding axes (30,31). Proposed statistical solutions, like the Tracy-Widom test, are similarly flawed for this purpose, as they identify significant axes of genetic variation, not necessarily axes correlated with confounding (21). The fundamental disconnect is that PCs are selected based on genetic variance, yet their utility in a full GRM model stems from their ability to partition genetic relatedness into components that can serve as proxies for social and environmental confounders. However, accurately modeling these complex ancestry-environment correlations is a major challenge. While some approaches attempt to infer environmental exposures from patterns of genetic relatedness (32), a more direct solution is to prioritize the collection of rich socio-demographic data. This would allow potential confounders to be modeled explicitly, rather than relying on PCs as imperfect proxies.

The development of scalable ancestral recombination graph (ARG) inference has enabled reframing existing methods but has not yet led to a new definition or measure of relatedness that accounts for the full genealogy. This work of reinterpretation is best exemplified by two recent, parallel developments. First, the GCTA-based GRM is now understood to be a statistical estimate of its true genealogical expectation, the eGRM, a specific application of the formal duality between genomic and genealogical statistics (33,34). Second, power of association testing has been boosted by replacing single variants with genealogical branches, using the ARG to test for latent variation (35). In these cases, the ARG has been used to provide a genealogical interpretation or re-framing of established statistical frameworks: the GRM and the single-locus association test, respectively. In contrast to reinterpreting existing methods, the GeSi framework is derived from the ground up by modeling the correlation of genetic effects given the coalescent times, and its resulting estimator, the cosine similarity of raw genotype vectors, is an emergent property, not a predefined inference target. Interestingly, however, the GeSi matrix converged to a standardized version of the GCTA/LDAK matrices (see genotype-based estimator in the **Suppl. Text**).

While our study robustly demonstrates the properties of GeSi and clarifies the role of different GRM types, its simulations have some limitations. The environmental confounder we modeled was a simplification of the true, unobserved social and environmental variables that influence complex traits. However, its purpose was conceptual: to demonstrate that when a known confounder correlated with population labels is explicitly included as a covariate, PCs are redundant for full GRMs. This isolates the effect of population structure and shows that it is not, by itself, the source of confounding that requires correction via PCs.

A key practical finding of our study was that GRMs calculated from WGS data yielded the most accurate heritability estimates, outperforming those from LD-pruned data, even when using the GCTA/LDAK GRMs that are typically calculated from LD-pruned data. We caution that this result may be partially attributed to our simulation design, which sampled causal variants uniformly along the genome. Our derivation of GeSi (see **Supplementary Text**) assumed that the number of causal mutations is linear with time, an assumption that holds under uniform sampling but may be violated in the human genome, where causal variants are known to be non-uniformly distributed and enriched in specific functional regions with distinct LD patterns and subject to different levels of natural selection (36,37). In such real-world scenarios, WGS data could lead to the over- or under-representation of heritability from certain genomic regions. Consequently, while LD-pruning can introduce its own biases, some form of LD-aware weighting or pruning would likely be necessary for both GeSi and other full GRMs in analyses of real data (38,39). This practical consideration, however, does not alter our fundamental conclusions that full GRMs contain more genealogical information than shallow GRMs, and that the primary role of PCs in a mixed model is not to model the genetic covariance but to serve as proxies for non-genetic confounders correlated with specific axes of variation.

The primary strength of this work lies in providing a new, theoretically grounded framework for measuring genetic relatedness. GeSi is derived as a continuous measure of genealogical similarity based on coalescent theory. Our results establish a clear and useful distinction between full GRMs, like GeSi and GCTA/LDAK, which capture the entire spectrum of relatedness, and shallow GRMs like PC-Relate, which isolate recent kinship. Finally, our study also shows that the role of principal components in mixed linear models is not to model genetic covariance due to population structure, but to function as proxies of confounders correlated with specific ancestry components represented as axes of variation in the PCA space.

Our framework opens several paths for future investigation. The unexpected finding that the standard GCTA method, which is an average of per-site ratios, also performs well with WGS data requires further exploration. Future simulations should explore the robustness of both GeSi and GCTA to this finding under different genetic architectures where causal variants are not sampled uniformly but are instead enriched in regions of high or low LD. Furthermore, the genealogical basis of GeSi makes it a promising tool for partitioning heritability into components explained by variants across different allele age bins, providing a more detailed understanding of a trait’s genetic architecture.

## 5. Conclusions

We introduce the coefficient of genealogical similarity (GeSi), a metric derived from coalescent theory that measures genetic relatedness as a continuum, unifying both familial relationships and shared ancestry. Our analyses show that this approach leads to a distinction between genetic relationship matrices based on the scope of the information they contain. Full GRMs, including GeSi, GCTA, and LDAK, retain the complete genealogical history of samples. In contrast, shallow GRMs, such as PC-Relate, are designed to discard deep ancestral information. This framework clarifies that the well-documented bias of methods like GCTA in structured populations is not a statistical artifact, but a direct consequence of capturing the full spectrum of relatedness. The practical benefit of retaining this information is demonstrated by more accurate heritability estimates and superior prediction of an individual’s total genetic value.

This work also redefines the role of principal components (PCs) in mixed models. We demonstrate that when a full GRM is used, PCs are redundant for modeling genetic covariance due to population structure. Their demonstrated utility in real-world analyses does not arise from modeling genetic effects, but from serving as proxies for unmeasured, non-genetic confounders correlated with ancestry. Our findings directly challenge two widespread practices in the field: the arbitrary partitioning of genetic relatedness into recent and distant components, and the conflation of population structure with confounding. Population structure is a source of genetic covariance that a full GRM correctly models; true confounding arises from external factors that correlate with that genetic structure. This study provides a new, theoretically-grounded framework for measuring relatedness and calls for a more precise, mechanistically-aware correction of confounding effects in genetics.

## 6. Acknowledgments

We are grateful to Sebastian Zollner, Eric Kernfeld, Jeff O’Connel, Braxton Mitchell, and Bing Guo. Their insightful comments and critical feedback during productive discussions were invaluable in refining the analyses and arguments presented in this manuscript.

