## Supplementary Figures 1-15 for "Unifying population structure and relatedness analysis through a coalescent approach"

### List of Supplementary figures


- **Supplementary figure 1**. Accuracy of true causal effects estimation in single-variant mixed linear model association tests in the Msprime-generated Simulation Set #2 ($\alpha$ =0).
- **Supplementary figure 2**. Accuracy of true causal effects estimation in single-variant mixed linear model association tests in the Msprime-generated Simulation Set #3 ($\alpha$ =−1).
- **Supplementary figure 3**. Type I error rate and inflation of the chi-square statistic in Simulated Set #2 (LDAK's $\alpha$=0).
- **Supplementary figure 4**. Type I error rate and inflation of the chi-square statistic in Simulated Set #3 (LDAK's $\alpha$=−1).
- **Supplementary figure 5**. Statistical power to detect a true causal variant in simulated set #2 ($\alpha$=0).
- **Supplementary figure 6**. Statistical power to detect a true causal variant in simulated set #3 ($\alpha$=−1).
- **Supplementary Figure 7**. Squared error of the heritability estimates under different levels of recent relatedness and admixture when the true scaling factor is $\alpha$=0 in the Simulation Set #2.
- **Supplementary Figure 8**. Distribution of heritability estimates under different levels of recent relatedness and admixture when the true scaling factor is $\alpha$=0 corresponding to the squared errors shown in Supplementary Figure 7.
- **Supplementary Figure 9**. Distribution of heritability estimates under different levels of recent relatedness and admixture when the true scaling factor is $\alpha$=−1, corresponding to the squared errors shown in Figure 2.
- **Supplementary Figure 10**. Square error and heritability estimate in simulations with and without confounding.
- **Supplementary figure 11**. Accuracy of the height prediction via BLUP in real data from the BioMe cohort.
- **Supplementary Figure 12**. Effect of the alpha parameter on the squared error of the heritability estimates.
- **Supplementary figure 13**. Selection of the alpha parameter for heritability estimation of human height.
- **Supplementary figure 14**. Group-specific heritability estimates of human height based on the $\alpha$ value that minimizes the AIC.
- **Supplementary Figure 15**. Group-specific heritability estimates of human height keeping a fixed at -1.

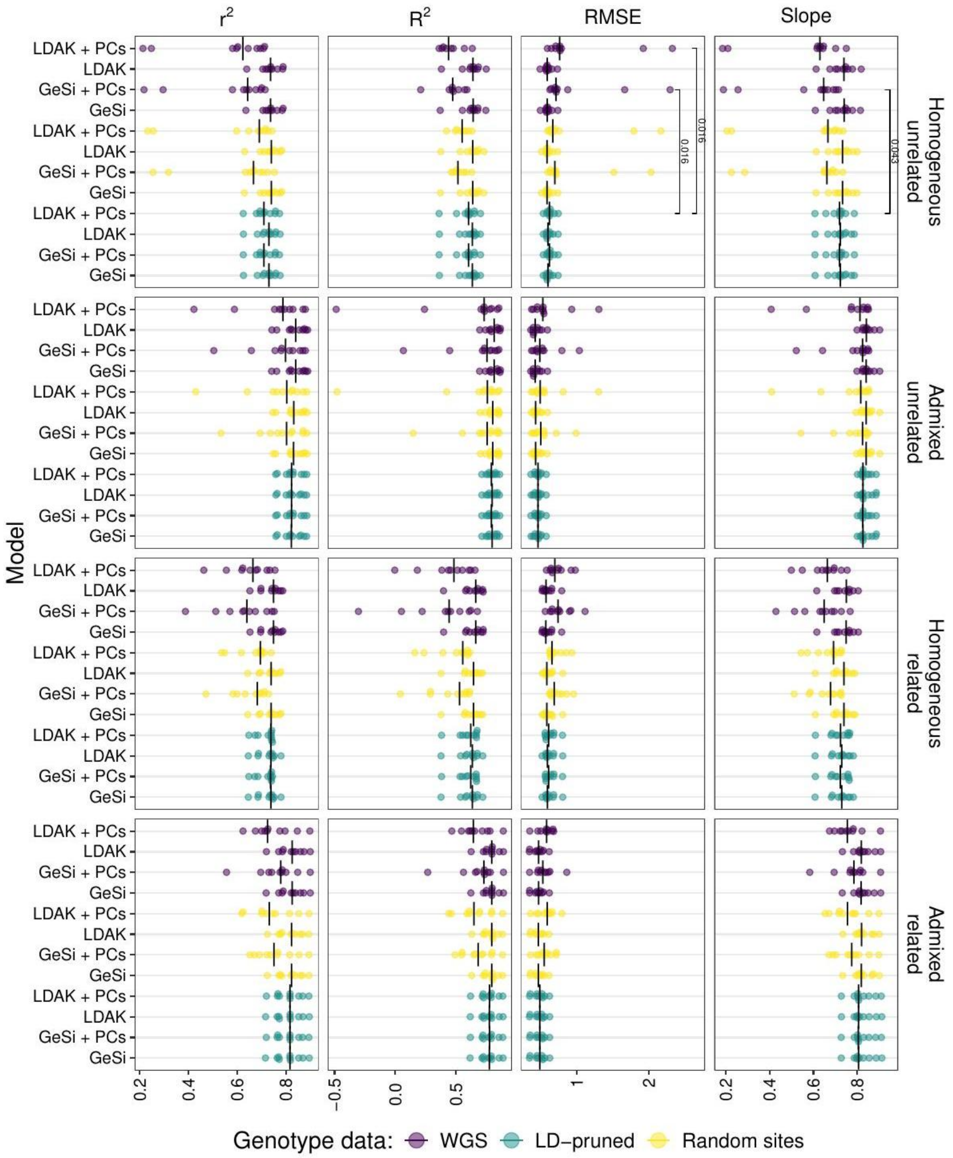


**Supplementary figure 1**. Accuracy of true causal effects estimation in single-variant mixed linear model association tests in the Msprime-generated Simulation Set #2 ($\alpha = 0$). Each column shows a different accuracy measure of the estimated SNP effect sizes: squared Pearson correlation coefficient ($r^2$), coefficient of determination ($R^2$), root of the mean squared error (RMSE), and the slope of regressing the effect size estimates on the true effect sizes (Slope). The black vertical bars represent the median for each measure within each panel. Each panel represents one of the four different combinations of demographic and ascertainment scenarios that were simulated: A) Unrelated individuals from Iberians in Spain (IBS); B) Mixture of related and unrelated individuals from IBS; C) Unrelated individuals from Latino and Latin American populations (LAT; see main text and methods); and D) Mixture of unrelated and unrelated individuals from LAT. Within each panel, the vertical axis details whether the mixed linear model (MLM) used a GeSi or LDAK genetic relationship matrix, and whether principal components (PCs) were included in the model. Ten replicates are shown for each model. Each GRM was calculated from three different types of color-coded genetic data: Whole Genome Sequence (WGS) data, LD-pruned data, or randomly selected sites. A two-sided Wilcoxon rank sum test was used to compare, within each panel, all methods against the reference model (LDAK-LD-pruned + 10 PCs). Only the FDR-adjusted p-values below 0.05 are shown.

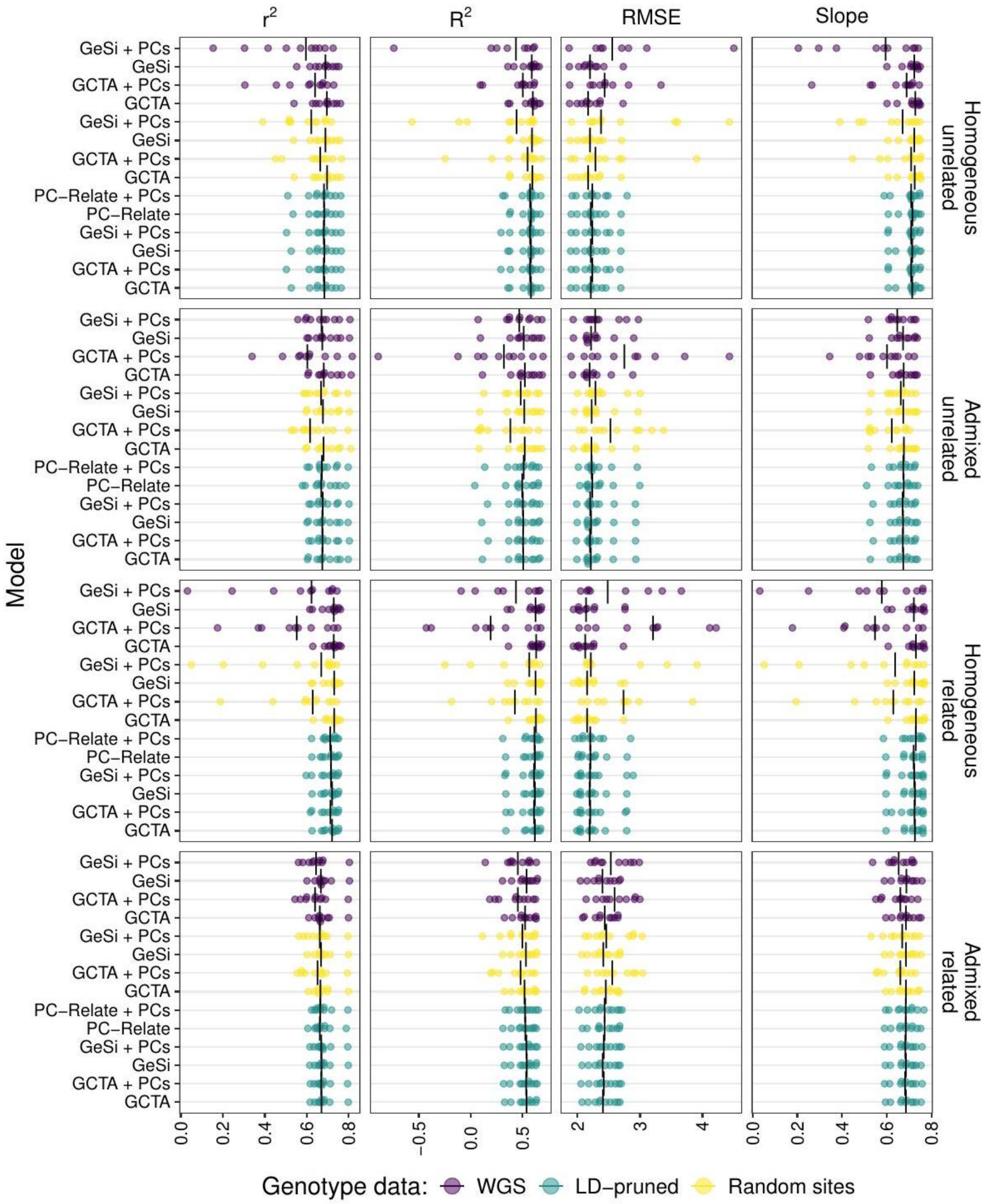


**Supplementary figure 2**. Accuracy of true causal effects estimation in single-variant mixed linear model association tests in the Msprime-generated Simulation Set #3 ($\alpha = -1$). Similar to Suppl. Fig. 1, but for the Simulation Set #3 ($\alpha = -1$). Note that the GCTA GRM is equivalent to LDAK with $\alpha = -1$.

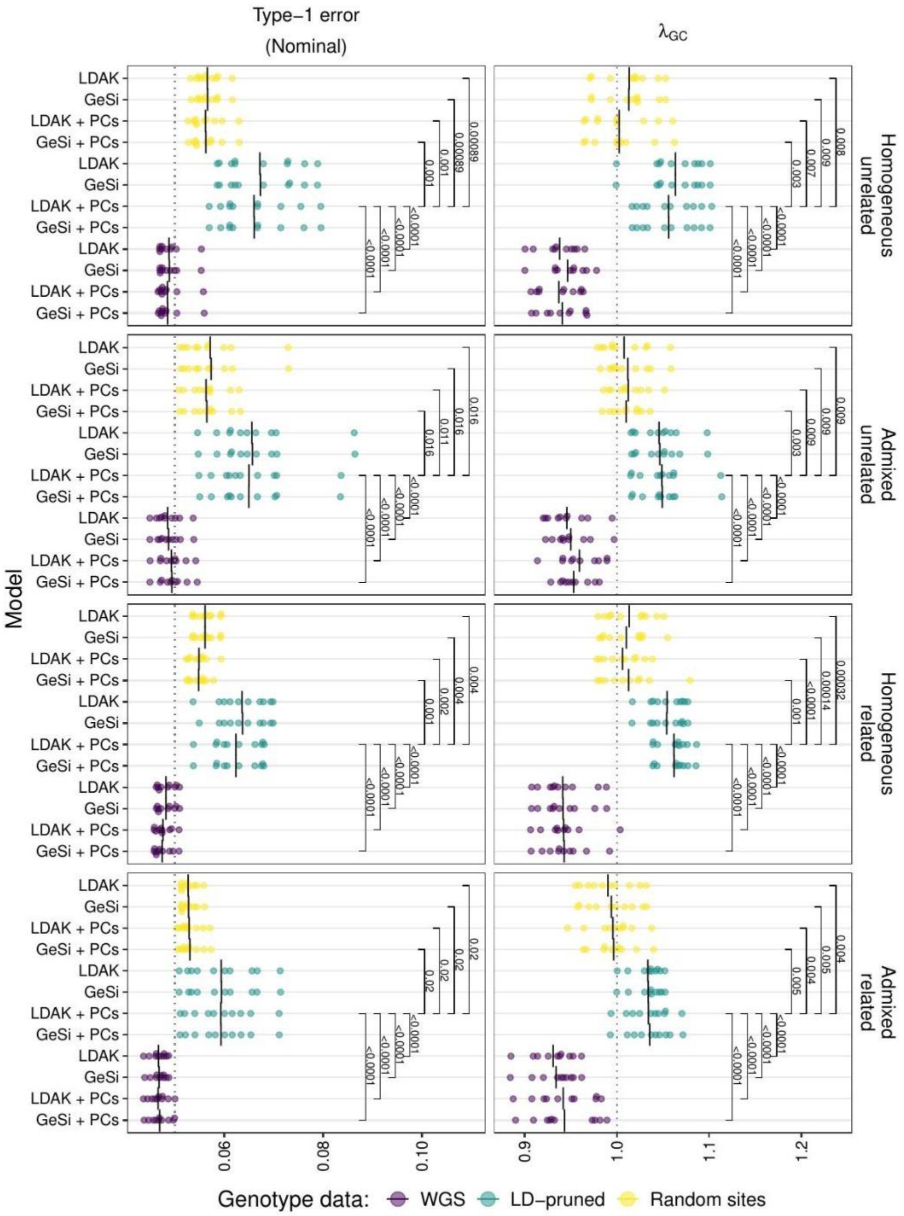


**Supplementary figure 3**. Type I error rate and inflation of the chi-square statistic in Simulated Set #2 (LDAK's $\alpha = 0$). The first column shows the type I error rate at a significance threshold of 0.05; and the second column shows the genomic control lambda parameter, defined as the median of the chi-square statistic divided by the median of a chi-square distribution with one degree of freedom. The black vertical bars show the mean for each measure within each panel. Each row represents one of the four different combinations of demographic and ascertainment scenarios that were simulated: A) Unrelated individuals from Iberians in Spain (IBS); B) Mixture of related and unrelated individuals from IBS; C) Unrelated individuals from Latino and Latin American populations (LAT; see main text and methods); and D) Mixture of unrelated and unrelated individuals from LAT. Within each panel, the vertical axis details whether the mixed linear model (MLM) used a GeSi or LDAK genetic relationship matrix, and whether principal components (PCs) were included in the model. Ten replicates are shown for each model. Each GRM was calculated from three different types of color-coded genetic data: Whole Genome Sequence (WGS) data, LD-pruned data, or randomly selected sites. A two-sided Wilcoxon rank sum test was used to compare, within each panel, all methods against the reference model (LDAK-LD-pruned + 10 PCs). Only the FDR-adjusted p-values below 0.05 are shown.

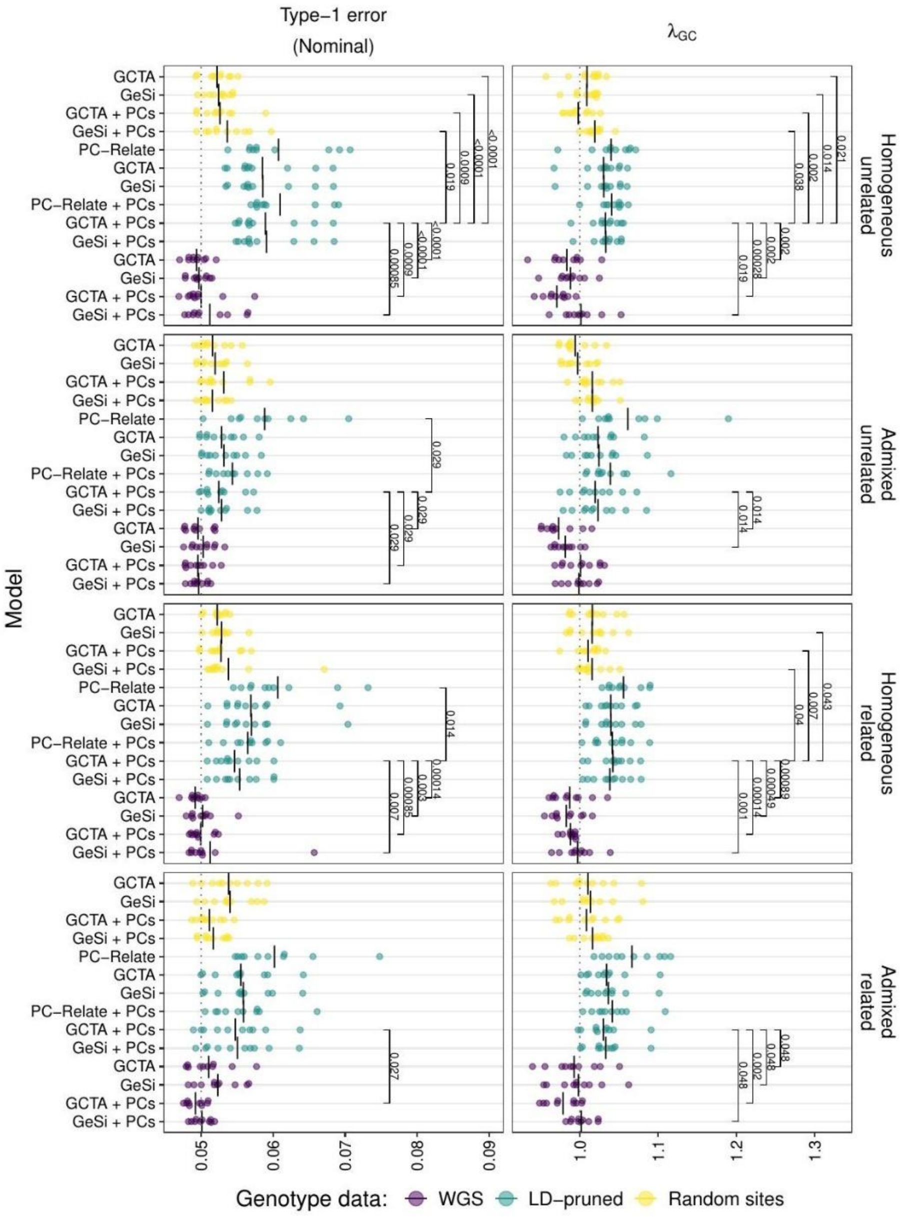


**Supplementary figure 4**. Type I error rate and inflation of the chi-square statistic in Simulated Set #3 (LDAK's $\alpha = -1$). Similar to Suppl. Fig. 3, but for the Simulation Set #3 ($\alpha = -1$). Note that the GCTA GRM is equivalent to LDAK with $\alpha = -1$.

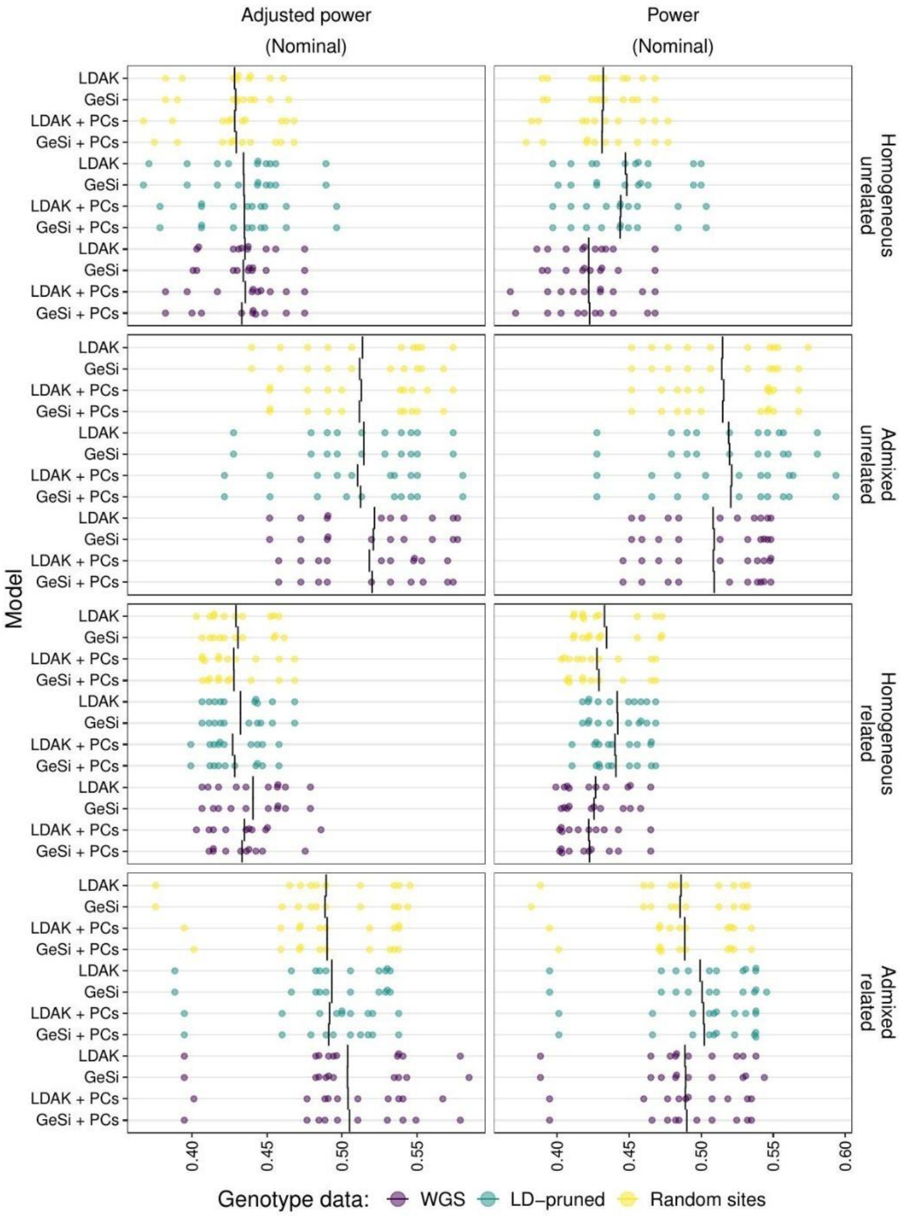


**Supplementary figure 5**. Statistical power to detect a true causal variant in simulated set #2 ($\alpha = 0$). The first column shows the $\lambda_{GC}$-adjusted power to detect a true causal variant, and the second column shows the statistical power without adjusting for inflation. Each row represents one of the four different combinations of demographic and ascertainment scenarios that were simulated: A) Unrelated individuals from Iberians in Spain (IBS); B) Mixture of related and unrelated individuals from IBS; C) Unrelated individuals from Latino and Latin American populations (LAT; see main text and methods); and D) Mixture of unrelated and unrelated individuals from LAT. Within each panel, the vertical axis details whether the mixed linear model (MLM) used a GeSi or LDAK genetic relationship matrix, and whether principal components (PCs) were included in the model. Ten replicates are shown for each model. Each GRM was calculated from three different types of color-coded genetic data: Whole Genome Sequence (WGS) data, LD-pruned data, or randomly selected sites. A two-sided Wilcoxon rank sum test was used to compare, within each panel, all methods against the reference model (LDAK-LD-pruned + 10 PCs). However, none of the comparisons was significant and therefore are not shown (FDR-adjusted p-values > 0.05).

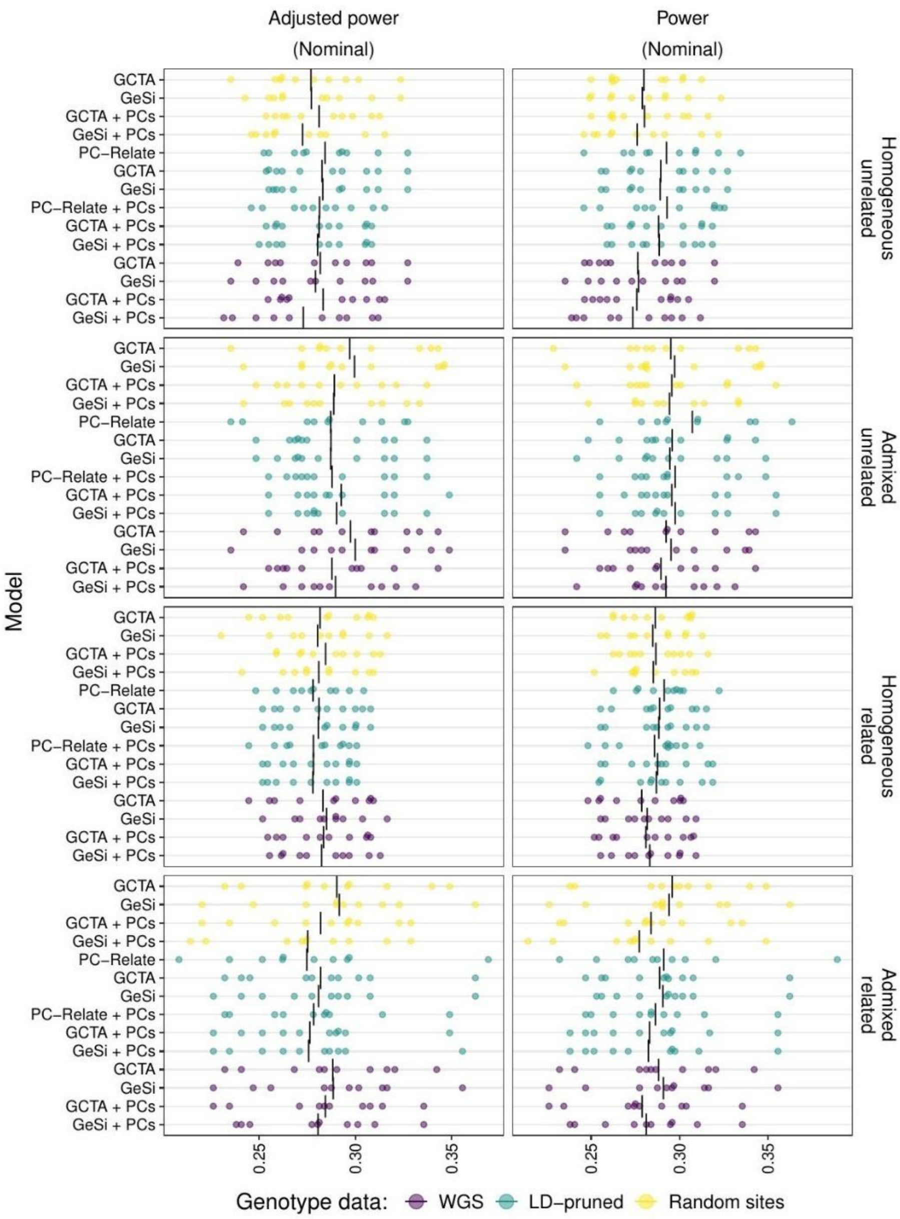


**Supplementary figure 6**. Statistical power to detect a true causal variant in simulated set #3 ($\alpha = -1$). Similar to Suppl. Fig. 5, but for the Simulation Set #3 ($\alpha = -1$). Note that the GCTA GRM is equivalent to LDAK with $\alpha = -1$.

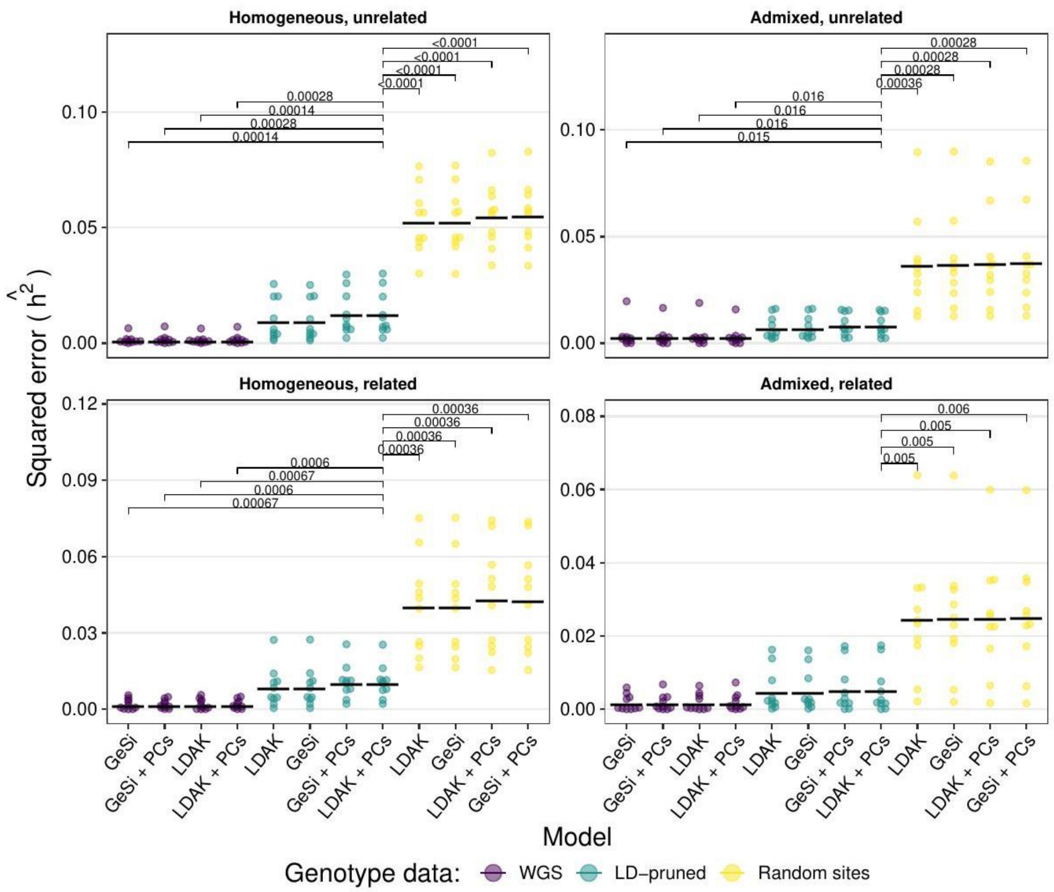


**Supplementary Figure 7**. **Squared error of the heritability estimates under different levels of recent relatedness and admixture when the true scaling factor is $\alpha = 0$ in the Simulation Set #2**. The X axis details whether the mixed linear model (MLM used a GeSi or LDAK genetic relationship matrix, and whether principal components (PCs) were included in the model. Ten replicates are shown for each model. Each GRM was calculated from three different types of color-coded genetic data: Whole Genome Sequence (WGS) data, LD-pruned data, or randomly selected sites. Each panel represents one of the four different combinations of demographic and ascertainment scenarios that were simulated: A) Unrelated individuals from Iberians in Spain (IBS); B) Mixture of related and unrelated individuals from IBS; C) Unrelated individuals from Latino and Latin American populations (LAT; see main text and methods); and D) Mixture of unrelated and unrelated individuals from LAT. A two-sided Wilcoxon rank sum test was used to compare all methods against the MLM model with 10 PCs and an LDAK matrix calculated from LD-pruned data. Models are displayed from best to worst within each panel as judged based on the mean squared error. A square bracket and FDR-adjusted p-value are shown for all comparisons with an FDR <0.05. Black horizontal lines: Mean squared error.

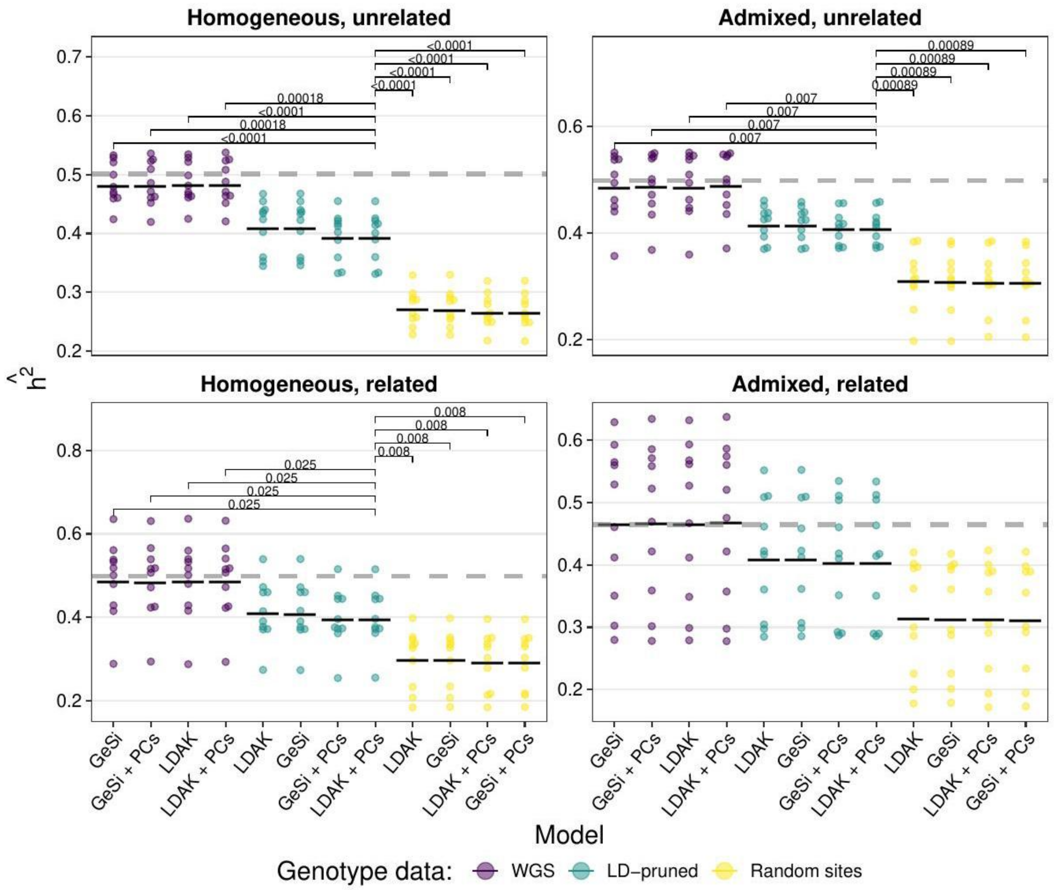


**Supplementary Figure 8**. Distribution of heritability estimates under different levels of recent relatedness and admixture when the true scaling factor is $\alpha = 0$, corresponding to the squared errors shown in Supplementary Figure 7. Heritability estimates correspond to the mean-square errors shown in Figure 2. Black horizontal lines: Mean heritability estimate.

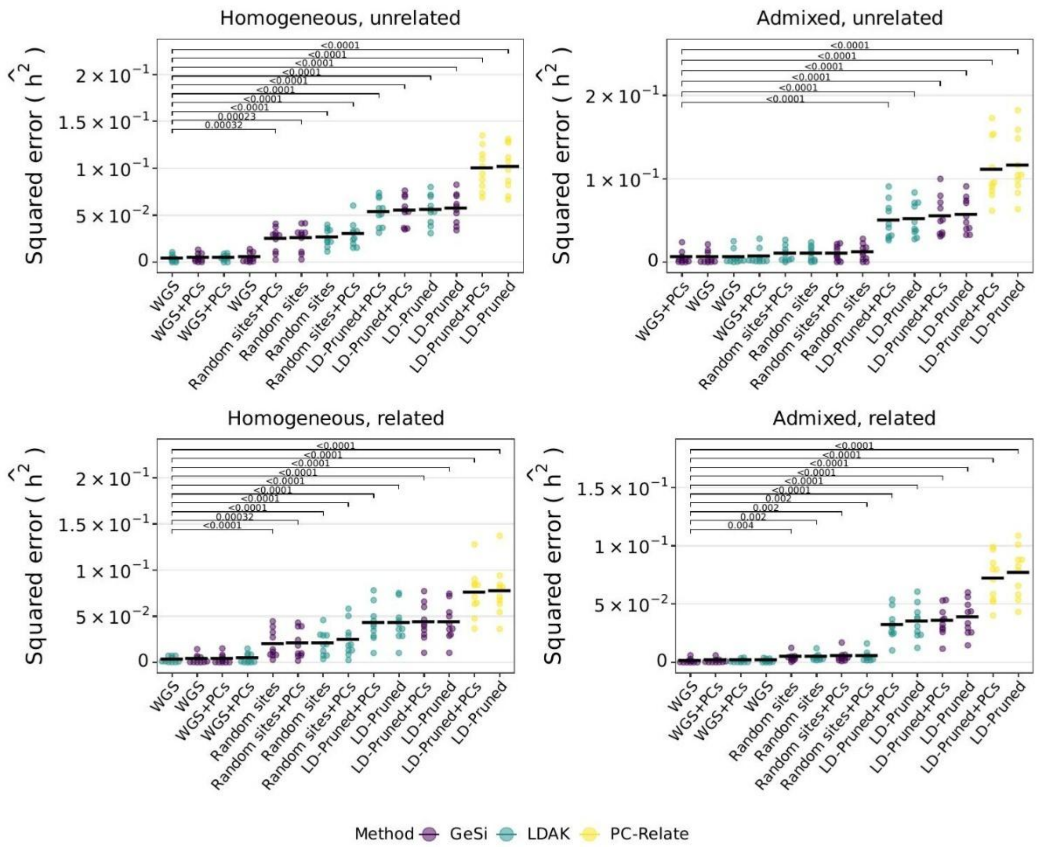


**Supplementary Figure 9**. Distribution of heritability estimates under different levels of recent relatedness and admixture when the true scaling factor is $\alpha = -1$, corresponding to the squared errors shown in Figure 2. Heritability estimates correspond to the mean-square errors shown in Figure 2. Black horizontal lines: Mean heritability estimate.

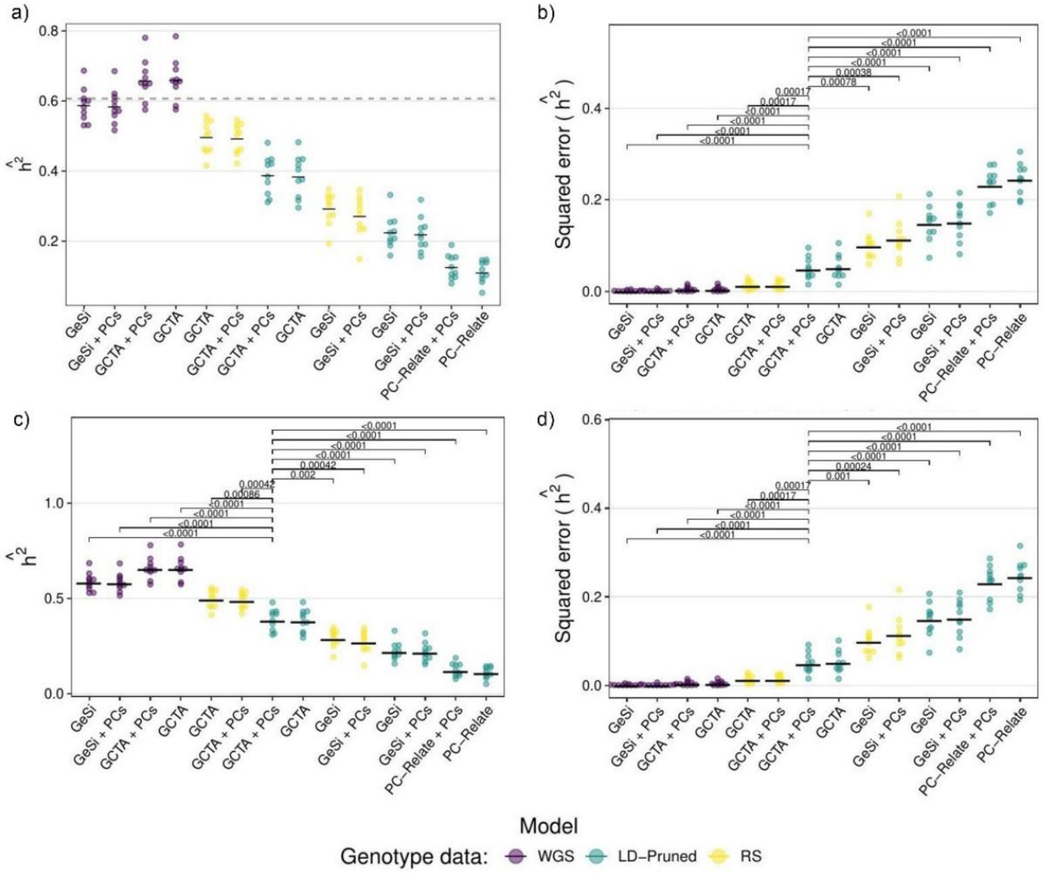


**Supplementary Figure 10. Square error and heritability estimate in simulations with and without confounding.** Within each panel, the horizontal axis details whether the mixed linear model (MLM) used a GeSi, LDAK or PC-Relate genetic relationship matrix, and whether principal components (PCs) were included in the model. All GRMs were calculated with $\alpha = -1$. Ten replicates are shown for each model. Each GRM was calculated from three different types of color-coded genetic data: Whole Genome Sequence (WGS) data, LD-pruned data, or randomly selected sites. Panels **a** and **b**: Simulation set #4 (no confounding). Panels **c** and **d**: Simulation Set #5 (confounding determined by demographic labels). A two-sided Wilcoxon rank sum test was used to compare all models against an MLM with 10 PCs and a GCTA matrix calculated from LD-pruned data. Models are displayed from best to worst within each panel as judged based on the mean squared error. A square bracket and FDR-adjusted p-value are shown for all comparisons with an FDR <0.05. Black horizontal lines: Mean-squared error

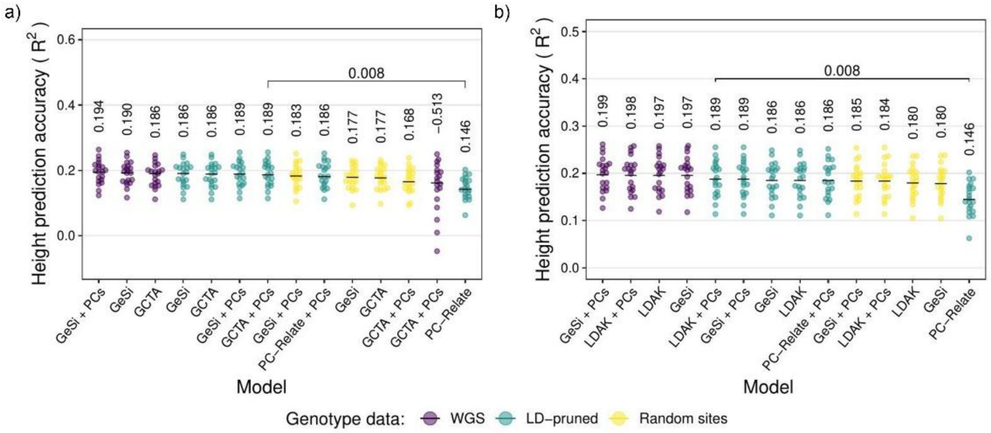


**Supplementary figure 11.** Accuracy of the height prediction via BLUP in real data from the BioMe cohort. **a)** Prediction accuracy keeping $\alpha$ fixed at -1. **b)** Prediction accuracy for the $\alpha$ value that maximizes the mean prediction accuracy. Each point represents the result of a cross-validation fold. The horizontal axis details whether the mixed linear model (MLM) used a GeSi, LDAK or PC-Relate genetic relationship matrix (GRM), and whether principal components (PCs) were included in the model. Each GRM was calculated from three different types of color-coded genetic data: Whole Genome Sequence (WGS) data, LD-pruned data, or randomly selected sites. A two-sided Wilcoxon rank sum test was used to compare all models against an MLM with 10 PCs and a GCTA matrix calculated from LD-pruned data. Models are displayed from best to worst within each panel as judged based on the mean squared error. A square bracket and FDR-adjusted p-value are shown for all comparisons with an FDR <0.05. Black horizontal lines: Mean-squared error.

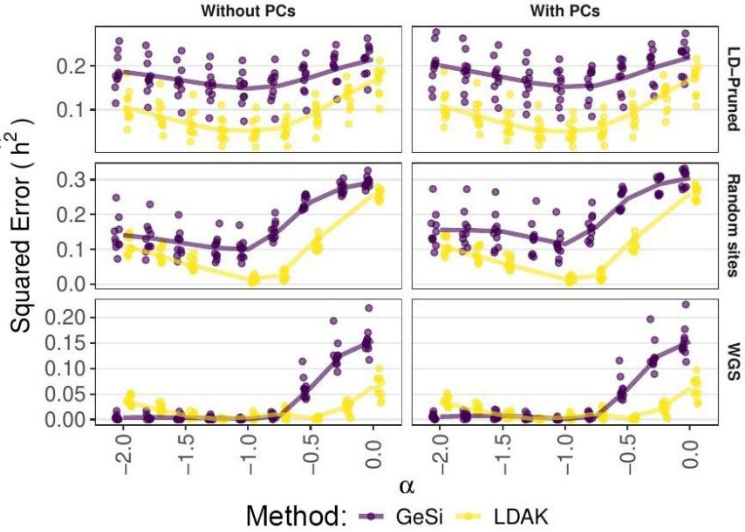


**Supplementary Figure 12**. Effect of the alpha parameter on the squared error of the heritability estimates. Each panel shows the heritability estimate squared error obtained from mixed linear models fit under different conditions specified by the facet titles. Each row indicates the type of genotype data (LD-pruned, Random sites or WGS) used to calculate the GRMs in the model. Each column indicates whether the model included principal components ("Without PCs" and "With PCs"). Each dot is a replicate (n=10), and the solid lines connect the mean of the ten replicates of each GRM method (purple for GeSi, yellow for LDAK). The GCTA is equivalent to the LDAK model with $\alpha = -1$. Data from Simulation Set #4 (no confounding, true alpha = -1).

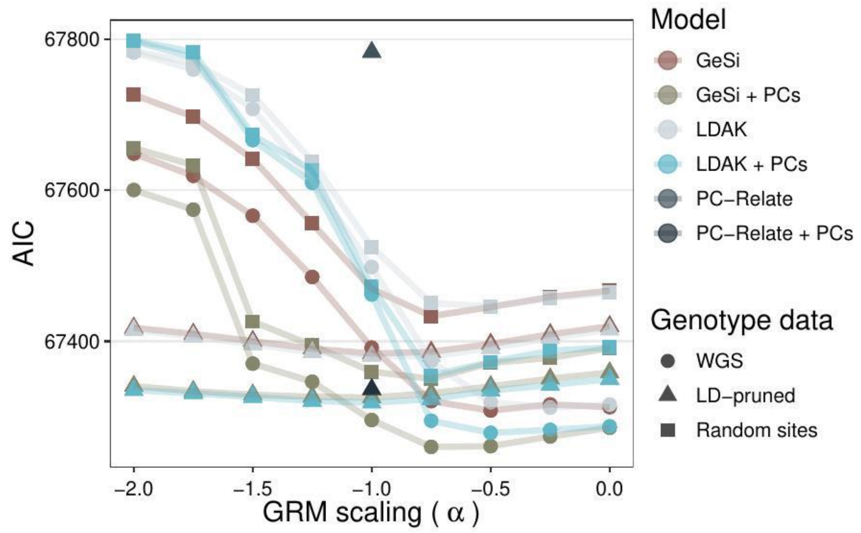


**Supplementary figure 13**. Selection of the alpha parameter for heritability estimation of human height. We defined six different lineal models based on the type of GRM they used (GeSi, LDAK/GCTA or PC-Relate) and on whether they used principal components as covariates or not. The GRM was calculated from the three different types of genotype data specified in the right-hand legend (WGS data, LD-pruned data or random sites), and with different $\alpha$ values, specified by the horizontal axis, raging from -2 to 0 in increments of 0.25. We calculated the AIC for each of the mixed lineal models defined by the combination of all these parameters. For each type of model and genotype data, we identified the $\alpha$ value that minimized the AIC of the model (see **Fig. 4** in main text).

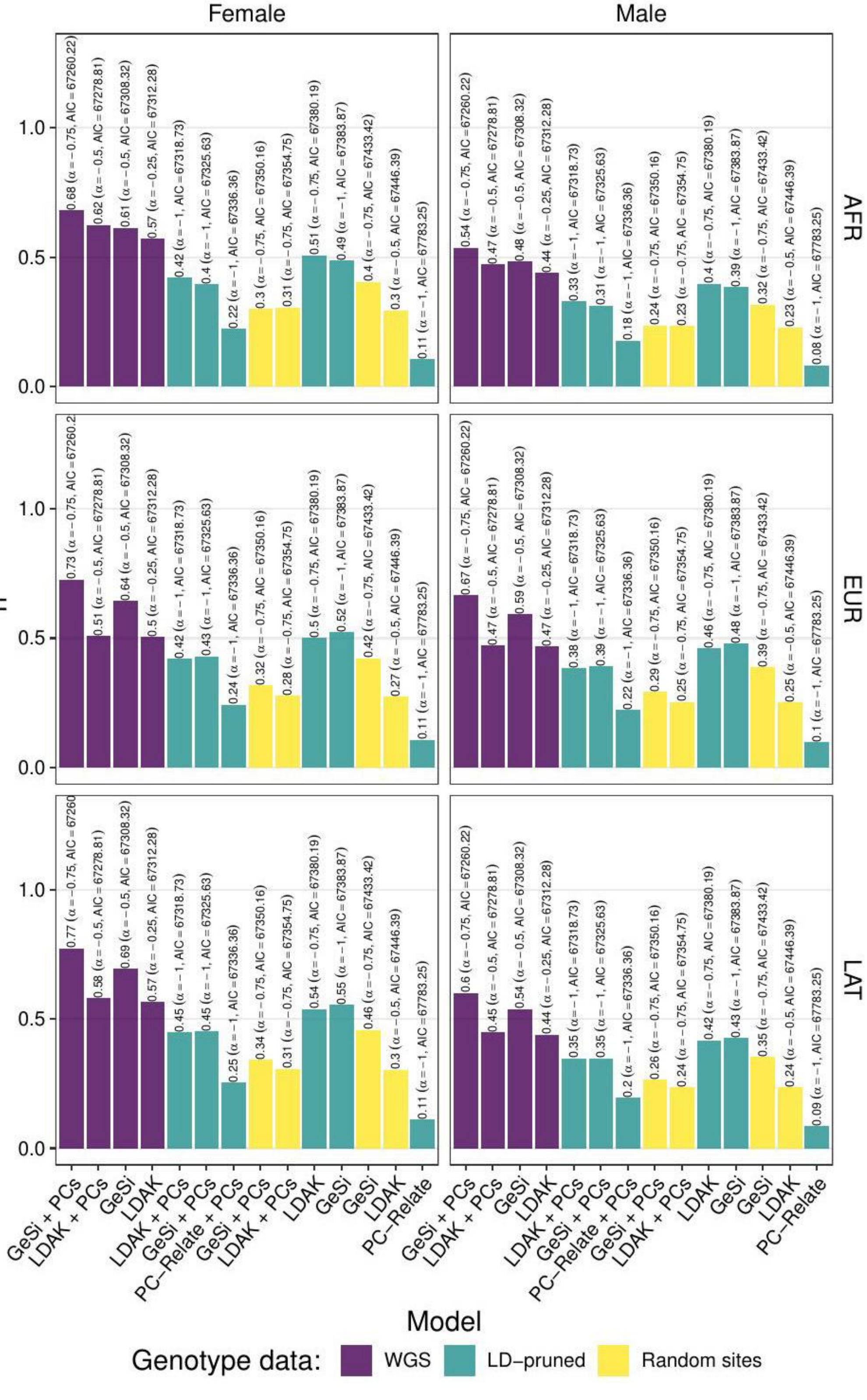

Female
Male
AFR
EUR
LAT
1.0
0.5
0.0
ĥ²
0.68 (α = −0.75, AIC = 67260.22)
0.62 (α = −0.5, AIC = 67278.81)
0.61 (α = −0.5, AIC = 67308.32)
0.57 (α = −0.25, AIC = 67312.28)
0.42 (α = −1, AIC = 67318.73)
0.4 (α = −1, AIC = 67325.63)
0.22 (α = −1, AIC = 67336.36)
0.3 (α = −0.75, AIC = 67350.16)
0.31 (α = −0.75, AIC = 67354.75)
0.51 (α = −0.75, AIC = 67380.19)
0.49 (α = −1, AIC = 67383.87)
0.4 (α = −0.75, AIC = 67433.42)
0.3 (α = −0.5, AIC = 67446.39)
0.11 (α = −1, AIC = 67783.25)
0.54 (α = −0.75, AIC = 67260.22)
0.47 (α = −0.5, AIC = 67278.81)
0.48 (α = −0.5, AIC = 67308.32)
0.44 (α = −0.25, AIC = 67312.28)
0.33 (α = −1, AIC = 67318.73)
0.31 (α = −1, AIC = 67325.63)
0.18 (α = −1, AIC = 67336.36)
0.24 (α = −0.75, AIC = 67350.16)
0.23 (α = −0.75, AIC = 67354.75)
0.4 (α = −0.75, AIC = 67380.19)
0.39 (α = −1, AIC = 67383.87)
0.32 (α = −0.75, AIC = 67433.42)
0.23 (α = −0.5, AIC = 67446.39)
0.08 (α = −1, AIC = 67783.25)
0.73 (α = −0.75, AIC = 67260.2
0.51 (α = −0.5, AIC = 67278.81)
0.64 (α = −0.5, AIC = 67308.32)
0.5 (α = −0.25, AIC = 67312.28)
0.42 (α = −1, AIC = 67318.73)
0.43 (α = −1, AIC = 67325.63)
0.24 (α = −1, AIC = 67336.36)
0.32 (α = −0.75, AIC = 67350.16)
0.28 (α = −0.75, AIC = 67354.75)
0.5 (α = −0.75, AIC = 67380.19)
0.52 (α = −1, AIC = 67383.87)
0.42 (α = −0.75, AIC = 67433.42)
0.27 (α = −0.5, AIC = 67446.39)
0.11 (α = −1, AIC = 67783.25)
0.67 (α = −0.75, AIC = 67260.22)
0.47 (α = −0.5, AIC = 67278.81)
0.59 (α = −0.5, AIC = 67308.32)
0.47 (α = −0.25, AIC = 67312.28)
0.38 (α = −1, AIC = 67318.73)
0.39 (α = −1, AIC = 67325.63)
0.22 (α = −1, AIC = 67336.36)
0.29 (α = −0.75, AIC = 67350.16)
0.25 (α = −0.75, AIC = 67354.75)
0.46 (α = −0.75, AIC = 67380.19)
0.48 (α = −1, AIC = 67383.87)
0.39 (α = −0.75, AIC = 67433.42)
0.25 (α = −0.5, AIC = 67446.39)
0.1 (α = −1, AIC = 67783.25)
0.77 (α = −0.75, AIC = 67260
0.58 (α = −0.5, AIC = 67278.81)
0.69 (α = −0.5, AIC = 67308.32)
0.57 (α = −0.25, AIC = 67312.28)
0.45 (α = −1, AIC = 67318.73)
0.45 (α = −1, AIC = 67325.63)
0.25 (α = −1, AIC = 67336.36)
0.34 (α = −0.75, AIC = 67350.16)
0.31 (α = −0.75, AIC = 67354.75)
0.54 (α = −0.75, AIC = 67380.19)
0.55 (α = −1, AIC = 67383.87)
0.46 (α = −0.75, AIC = 67433.42)
0.3 (α = −0.5, AIC = 67446.39)
0.11 (α = −1, AIC = 67783.25)
0.6 (α = −0.75, AIC = 67260.22)
0.45 (α = −0.5, AIC = 67278.81)
0.54 (α = −0.5, AIC = 67308.32)
0.44 (α = −0.25, AIC = 67312.28)
0.35 (α = −1, AIC = 67318.73)
0.35 (α = −1, AIC = 67325.63)
0.2 (α = −1, AIC = 67336.36)
0.26 (α = −0.75, AIC = 67350.16)
0.24 (α = −0.75, AIC = 67354.75)
0.42 (α = −0.75, AIC = 67380.19)
0.43 (α = −1, AIC = 67383.87)
0.35 (α = −0.75, AIC = 67433.42)
0.24 (α = −0.5, AIC = 67446.39)
0.09 (α = −1, AIC = 67783.25)
GeSi + PCs
LDAK + PCs
GeSi
LDAK
LDAK + PCs
GeSi + PCs
PC−Relate + PCs
GeSi + PCs
LDAK + PCs
LDAK
GeSi
GeSi
LDAK
PC−Relate
Model
Genotype data:
WGS
LD−pruned
Random sites

**Supplementary figure 14**. **Group-specific heritability estimates of human height based on the $\alpha$ value that minimizes the AIC.** We identified the value of the $\alpha$ parameter that minimized the AIC of each model defined on the horizontal axis and data type indicated in the legend. The models are sorted from lowest to highest AIC from left to right. We allowed for heterogeneous residual variance across each of the sex-by-population groups, and thus obtained six different heritability estimates for each model for each data type. The vertical axis shows the group-specific heritability estimates for each combination of sex and population labels (AFR, EUR, LAT) specified on the facets' titles. The selection of the $\alpha$ parameter value is detailed in **Suppl. Fig. 13**.

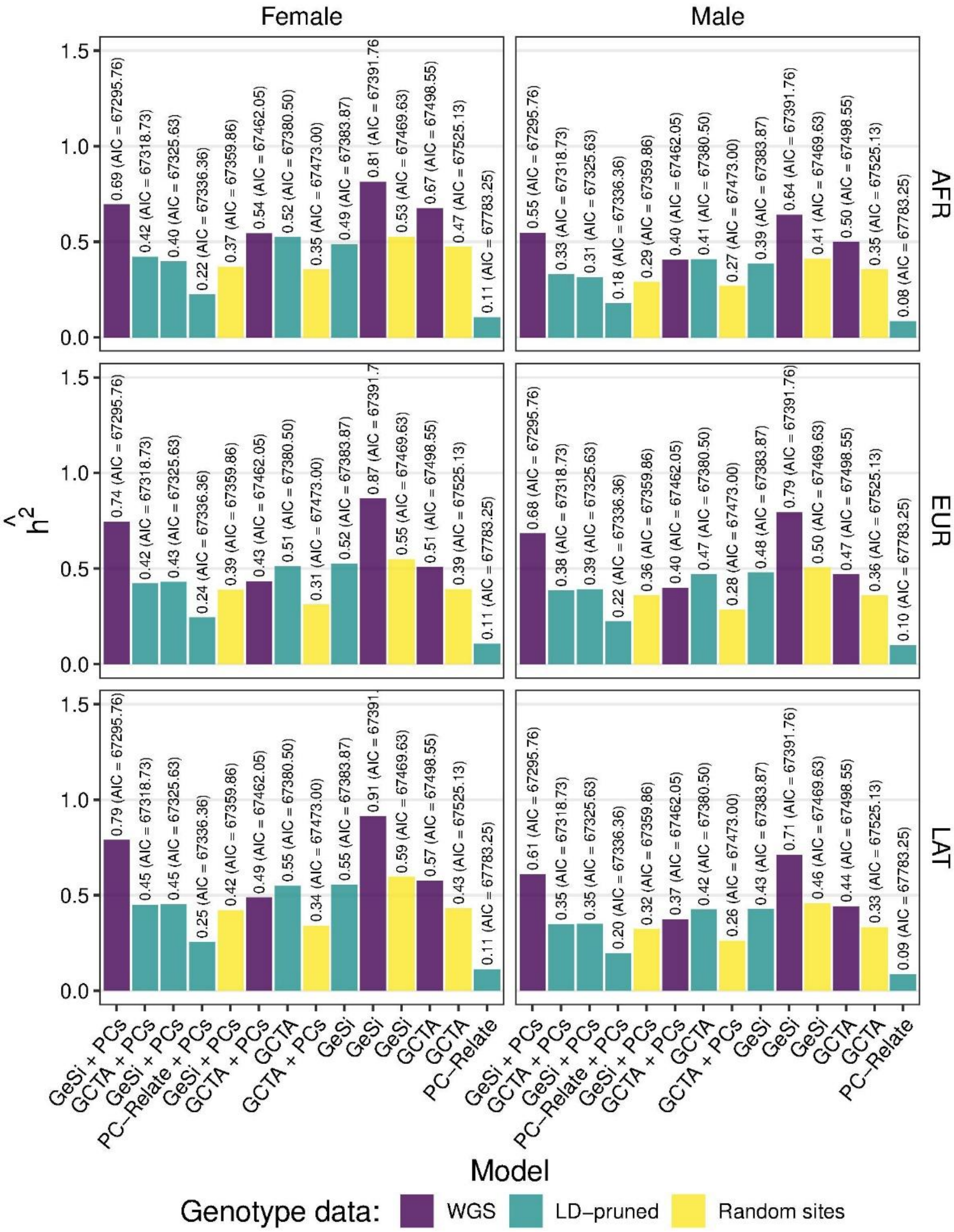


**Supplementary Figure 15. Group-specific heritability estimates of human height keeping $\alpha$ fixed at -1.** Similar to **Suppl. Fig. 14**, except that the value of the $\alpha$ parameter was fixed at -1.
