## Supplementary Text for "Unifying population structure and relatedness analysis through a coalescent approach"

### I. Methods

#### 1. Data sources and preparation

We used both real and simulated data. All simulated phenotypes were quantitative. The real phenotype data consisted of both quantitative (height and body mass index) and binary (coronary artery disease) traits. For each genotype dataset, we generated three variations: All biallelic single-nucleotide polymorphisms (SNPs) from whole-genome sequence (“WGS”), biallelic SNPs in linkage-equilibrium (“LD-pruned”), and randomly sampled SNPs from the WGS data (“RS”), with the number of sampled SNPs matching that of the LD-pruned data. All datasets included autosomal variants only. In this section, we describe the real-genotype datasets. The simulated datasets are described in a separate section.

##### 1.1. Peruvian Genome Project Plus (PGP+)

The Peruvian Genome Project Plus (PGP+, n = 1702) merged several datasets obtained though high-coverage whole-genome sequencing. The core dataset (PGP) included 147 Peruvians from the coastal (n=46), Andean (n=73) and Amazonian (n=28) regions. This core set of samples was expanded with a subset from the high-coverage releases of the 1000 Genomes Project[1] (1KGP, n = 1417), the Human Genome Diversity Project (HGDP, n = 117), and Simons Genome Diversity Project (SGDP, n = 21) in order to maximize the genetic diversity and the set of callable sites in the variant calling pipeline. The non-PGP set contained a total of 371 samples from Africa, 433 from the Americas, 334 from East Asia, 407 from Europe, and 10 from South Asia.

The HGDP FASTQ files were obtained from the BioProject PRJEB6463 at the NCBI’s Sequence Read Archive (SRA) using the fasterq-dump tool. Raw 30x WGS 1KGP FASTQ files were retrieved via IBM Aspera FASP from the ENA public run directories (/vol1/run/) corresponding to BioProject PRJEB31736/ERP114329.

We implemented a joint variant-calling pipeline for samples of all four sources. The FASTQ files were then quality-filtered using fastp[2] (version 0.23.2) in default mode to remove low-quality bases, short reads, and reads with too many uncalled bases (Ns). Filtered reads were then aligned to the GRCh38 reference genome using bwa-mem[3] (version 0.7.17-r1188) with read group header line information incorporated via the bwa -R parameter. For samples with multiple FASTQ sets due to multiple sequencing runs, we merged the aligned BAM files using samtools[4] merge (version 1.11) before further processing.

Joint genotyping was performed using an adapted pipeline based on the UK Biobank[5] and NYGC[1] workflows. We conducted base quality score recalibration (BQSR) using GATK with recognized known sites from dbSNP v.138 and Mills Gold Standard[6] mapped to the GRCh38 reference human genome downloaded from the Broad Institute’s public GATK resource bundle[7]. We used Picard[8] (v.2.25.3) to ensure the correct association of paired-end reads, and used samtools for sorting and marking duplicates. GATK's (v.4.2.6.1) HaplotypeCaller was run in GVCF mode for each sample, producing a single gVCF with genotype likelihood information for every site. We used GATK’s GenomicsDBImport and GenotypeGVCFs, splitting each chromosome into 2Mb intervals (keeping shorter segments at chromosome ends), with 1000 bp of interval padding. Genotyping was done in batches of 50 individuals. Failed interval runs were retried by increasing memory or using a different GATK version, following UK Biobank WGS pipeline recommendations.

For variant recalibration, we used GATK VariantRecalibrator and ApplyVQSR with dbSNP, HapMap, and 1KGP high-confidence files for SNPs and Mills and dbSNP files for INDELs. We applied truth sensitivity thresholds of 99.8 for SNPs and 99 for INDELs. Finally, we used bcftools[4] (v.1.15.1) to keep only PASS variants and to convert genotypes with depth (DP) < 10 and genotype quality (GQ) < 20 to missing data.

##### 1.2. BioMe cohort

The BioMe dataset originated from the BioMe cohort (dbGAP study accession: phs001644) and had phenotype and whole-genome sequence (WGS) data available for 12,054 subjects. We excluded samples with missing data for any of the variables: age, sex, height, body-mass index, coronary artery disease status, and peripheral artery disease status. We filtered out subjects missing basic covariates (population label, sex and age) or height data, potential duplicate entries. Finally, we kept only subjects described as African or African Americans (AFR, n= 2,975), European Americans (EUR, n = 2,442), and Latino or Latin American (LAT, n=4,468), making a total of 9,885 subjects in the final dataset. Subjects described as East Asians, South Asians and Indigenous Americans were filtered out due to the small sample sizes of those groups. Using PC-Relate (see methods) and a kinship coefficient threshold of $(1/2)^{4.5}$, corresponding to the lower-boundary of third-degree relatives[9], we identified a set of 9,190 unrelated subjects and 695 subjects related to them, which were described as AFR (n=206), EUR (n=28) and LAT (n=461). The WGS data was extracted from the TOPMed project Freeze 9 dataset[10].

#### 2. Phenotype simulation

We simulated traits with a number of different genetic architectures. Here we described the different strategies we used to simulate phenotypes. In section 3, we specify which phenotype simulation strategy was used for each dataset.

The genetic architecture was defined by the heritability, number of causal sites, dependence of the effect size on the minor allele frequency (LDAK's $\alpha$ parameter[11]), effect of the background linkage disequilibrium on the distribution of causal SNPs, and environmental confounding correlated with ancestry. All phenotype simulations were done in R v.4.3.3.

We simulated quantitative phenotypes under a polygenic additive genetic model. For each simulation, we selected at random a predefined number of causal biallelic SNPs (100, 1,000, or 10,000). The causal SNPs were sampled either with uniform probability, or sampling weight depending on their LD scores, as will be explained later. The effect sizes were drawn from a normal distribution with mean 0 and unit variance and were then scaled according to their minor allele frequency (MAF) via the $\alpha$ parameter as described in the next section.

For individual $i$, the total genetic effects were calculated as $g_i = \sum_j X_{ij}\beta_j$, where $X_{ij}$ represents the genotype for individual $i$ at causal SNP $j$, and $\beta_j$ is the effect size for causal SNP $j$. We set the narrow-sense heritability ($h^2$) to predetermined values (0.2, 0.5, or 0.8) by adding an environmental component $e_i$ drawn from a normal distribution with mean zero and variance adjusted to achieve the target heritability:

$$\sigma_e^2 = \frac{\sigma_a^2(1 - h^2)}{h^2}$$

The final phenotype was calculated as $y_i = g_i + e_i$.

##### 2.1. Effect of the minor allele frequencies on SNP effect sizes (α parameter)

We used Doug Speed's LDAK[11] approach to allow SNP effect sizes to vary with allele frequency. Let $\mathbf{X} \in \mathbb{R}^{n \times p}$ denote the unscaled genotype matrix of $n$ subjects at $p$ sites and let $x_{ij}$ denote the genotype of subject $i$ at site $j$, where $x_{ij} \in \{0,1,2\}$ denotes the number of copies of the alternate allele. Let $f_j$ denote the alternate allele frequency at site $j$, then the matrix of mean-centered genotypes $\mathbf{X}^{(c)}$ has elements $x_{ij}^{(c)} = \left(x_{ij} - 2f_j\right)$, and the scaled genotypes matrix $\mathbf{Z} \in \mathbb{R}^{n \times p}$ has elements $z_{ij} = x_{ij}^{(c)} \times [2f(1-f)]^{-\alpha/2}$. Let $\beta_j$ denote the effect size of variant $j$ on a trait in the raw genotype scale. Then, if the effect sizes have constant variance in the scaled genotype space, the variance of the raw effect sizes is $Var\left(\beta_j\right) = [2f(1-f)]^{\alpha} \times constant$, and $E[h^2] \propto [2f(1-f)]^{1+\alpha}$ is the expected heritability explained by site $j$. The LDAK model reduces to the GCTA model by setting $\alpha = -1$.

To use the LDAK model, we first simulated the effects sizes $\beta_j^{(raw)}$ from a standard normal distribution and assigned them to the randomly selected sites. Then we obtained the final effect sizes with $\beta_j = \beta_j^{(raw)} \times \left[2f_j\left(1-f_j\right)\right]^{-\alpha/2}$.

#### 2.2. Simulation of confounding effects

The confounding effects were simulated to correlate with population structure, specifically related to the three population labels European Americans (EUR), African or African Americans (AFR), and Latino or Latin Americans (LAT). An incidence matrix $\mathbf{Q} \in \mathbb{R}^{n \times 3}$ was constructed, mapping each of the $n$ subjects to their respective population. The vector $\mathbf{b}^T = [b_{EUR}, b_{LAT}, b_{AFR}]$ contains the effect sizes associated with each population. Then, we defined the model $\mathbf{Y} = \mathbf{X\beta} + \mathbf{Qb} + \mathbf{e}$, where $\mathbf{X}$ is the genotype matrix of causal sites and $\boldsymbol{\beta}$ a vector of their coefficients.

We followed the next procedure to introduce a specific directional confounding pattern that correlated with the amount of both European and African ancestry:

1. The effect $b_{raw(LAT)} = b_{(LAT)}$ was set to zero, $b_{raw(EUR)}$ was sampled from a truncated negative standard normal distribution, and $b_{raw(AFR)}$ from a truncated positive standard normal distribution. Then we defined the vector $\mathbf{b}_{raw}^T = [b_{raw(EUR)}, b_{raw(LAT)}, b_{raw(AFR)}]$. We set $b_{(LAT)}$ to zero because LAT had intermediate levels of both European and African ancestry compared to the groups labeled as EUR and AFR.
2. A scaling factor $\omega$ was calculated to ensure that the genetic and confounding effects explained the same amount of phenotypic variance. This factor was calculated by $\omega = \left(\frac{\mathrm{Var}(\mathbf{X\beta})}{\mathrm{Var}(\mathbf{Qb}_{raw})}\right)^{1/2}$. Where $\sigma_a^2 \coloneqq \widehat{\mathrm{Var}}(\mathbf{X\beta})$ is the variance of the total additive genetic effects, and $\mathrm{Var}(\mathbf{Qb}_{raw})$ is the variance contributed by the confounding effects using the raw coefficients $\mathbf{b}_{raw}$.
3. The final confounding effect vector $\mathbf{b}$ was obtained by multiplying the raw coefficient vector by the scaling factor: $\mathbf{b} = \omega\mathbf{b}_{raw}$.

This procedure ensured that the observed Pearson's correlation coefficient between genetic and confounding effects remained within the range of -0.18 to -0.09. The residual error term $\mathbf{e}$ was drawn independently from a normal distribution with mean zero and variance $\sigma_e^2$. The residual variance parameter was defined as $\sigma_e^2 = \frac{\sigma_a^2(1-h^2)}{h^2}$, where $h^2$ is the pre-specified narrow-sense heritability.

### 3. Simulation of genealogical trees and whole-genome sequence (WGS) data

We generated multiple simulated data sets to test different scenarios. Simulation Set #1 consisted of only genotype data; Sets #2 and #3 consisted of both simulated genotype and phenotype data; and Sets #4-#7 consisted of phenotypes simulated from real genotype data.

##### 3.1. Exploratory Simulation (Simulation Set #1)

We used Msprime[12,13] to simulate genealogical trees and whole-genome sequence data using a demographic model Latin American and reference continental ancestries[14]. Specifically, we used the backward-in-time Discrete-Time Wright-Fisher model[15] (DTWF) to generate data resembling eight populations: Chinese in Beijing (CHB), Colombians in Medellin (CLM), Iberians in Spain (IBS), Ancestral Indigenous Mexicans (MXB), Mexicans in Los Angeles (MXL), Peruvians in Lima (PEL), Puerto Ricans (PUR), and Yoruba in Ibadan, Nigeria (YRI). Each population consisted of 60 individuals, including 30 unrelated samples, five full-sib pairs, five half-sib pairs, and five first-cousin pairs. Three chromosomes (chr20-chr22) were simulated using the corresponding GRCh38 recombination maps, a mutation rate of $1.25x10^{-8}$ per site per generation, and the Jukes and Cantor (JC69) nucleotide substitution model[16]. Phenotypes were not simulated for this dataset.

##### 3.2. Latin American Demographic Model (Simulation Sets #2 and #3)

We modified the previously described demographic model of Latin American[14], by adding a new artificial population (LAT, short for Latino and Latin Americans ) resulting from an admixture event three generations ago consisting of 25% CLM, 25% MXL, 25% PEL, and 25% PUR. We set the admixture event three generations in the past so that current samples could draw their grandparents from any of the contributing populations. Similarly to Simulation Set #1, we used the DTWF and JC69 models, and mutation rate of $1.25x10^{-8}$, but generated ten chromosomes (chr13-chr22) using the GRCh38 recombination maps. We generated 5,000 samples under four different scenarios:

1. No recent admixture and no recent relatedness: 5,000 unrelated samples from IBS.
2. Admixture without recent relatedness: 5,000 unrelated samples from LAT.
3. No recent admixture with recent relatedness: 600 full-sib pairs, 600 half-sib pairs, 600 first-cousin pairs, and 1,400 unrelated subjects from IBS.
4. Admixture with recent relatedness: 600 full-sib pairs, 600 half-sib pairs, 600 first-cousin pairs, and 1,400 unrelated subjects from LAT.

For both Simulation Set #2 and Set #3, we simulated a polygenic trait with 0.50 narrow-sense heritability determined by 1,000 SNPs sampled at random with uniform probability. The SNP effect sizes were generated under the $GeSi_{tree}$ model ($\alpha = 0$, explained in the next section) in Simulation Set #2, and under the GCTA model for Simulation Set #3 ($\alpha = -1$). Both simulation sets consisted of 10 replicates per scenario.

##### 3.3. BioMe-Based Simulations (Simulation Sets #4 and #5)

We also ran simulations using real genotypes from the BioMe WGS data, assigning simulated phenotypes under various scenarios:

- **Simulation Set #4**: Phenotypes simulated with $\alpha = -1$, causal SNPs sampled with uniform probability and no confounding.

- **Simulation Set #5**: Phenotypes simulated with $\alpha = -1$, causal SNPs sampled with uniform probability, and environmental confounding weakly or moderately correlated with genetic ancestry.

We randomly selected 1,000 causal variants and set the heritability to 0.5 in the EUR group. All simulations were replicated 10 times.

#### 4. Derivation of the Coefficient of Genealogical Similarity (GeSi)

In this section, we use tree branch statistics to express the correlation in the number of derived alleles carried by two individuals. We then demonstrate that this branch statistics-based coefficient represents the pairwise phenotypic correlation between individuals under the assumption that $\alpha = 0$, i.e., that the causal variants' effect sizes are independent of their minor allele frequency. We then demonstrate that the pairwise cosine similarity of the genotype vectors is an estimator of this branch-based statistic, and thus of the Pearson correlation coefficient of the phenotype values.

##### 4.1. GeSi as the correlation of the number of derived alleles carried by two diploid individuals

To model the correlation between the number of mutations carried by two diploid individuals, we first model the covariance in the number of derived alleles carried by two haplotypes given t heir realized genealogy, and the variance in the number of mutations carried by any haplotype. We then use these components to model the correlation between the number of derived alleles carried by two diploid individuals.

##### 4.2. Covariance of the number of derived alleles carried by two haplotypes

Let $i$ and $j$ be two nodes representing the genealogy of any two haplotypes (see Fig. 1 in the main text), and let $N_i$, $N_j$ be two random variables (r.v.) representing the number of mutations they have accumulated since they diverged from the ancestral sequence represented by the root of the tree. Let $T_{ij}^{(s)}$ be the time between the root of the tree and the time when the genealogies of $i$ and $j$ diverged and let $T_{ij}^{(u)}$ be the time since their genealogies diverged until present time. Finally, let $T^{(R)}$ denote the time from present time to the root of the tree. We call $T_{i,j}^{(s)}$ the shared divergence time of haplotypes $i$, $j$, and $T_{ij}^{(u)}$ their unshared divergence time. Let $N_{i,j}^{(s)}$ be a r.v. representing the number of mutations that occurred during time $T_{ij}^{(s)}$, and $N_i^{(u)}$ and $N_j^{(u)}$ two r.v. representing the number of mutations that occurred during time $T_{ij}^{(u)}$ on the haplotypes represented by nodes $i$ and $j$. We define:

$$N_i = N_{ij}^{(s)} + N_i^{(u)}$$

$$N_j = N_{ij}^{(s)} + N_j^{(u)}$$

Then:

$$Cov(N_i, N_j) = Cov\left[\left(N_{ij}^{(s)} + N_i^{(u)}\right), \left(N_{ij}^{(s)} + N_j^{(u)}\right)\right]$$

We assume that the number of mutations arising on a haplotype follows a Poisson distribution given by the length $L$ of the sequence, the mutation rate $v$, and the divergence time. Thus:

$$N_i \sim Poisson(\lambda_{ij}^{(s)} + \lambda_{ij}^{(u)})$$

$$N_j \sim Poisson(\lambda_{ij}^{(s)} + \lambda_{ij}^{(u)})$$

Where $\lambda_{ij}^{(s)} = vLT_{ij}^{(s)}$ and $\lambda_{ij}^{(u)} = vLT_{ij}^{(u)}$

The number of mutations occurring on haplotypes $i$ and $j$ after their genealogies diverged are independent of each other. Thus, the covariance in the number of mutations carried by any two haplotypes is given by $Cov(N_i, N_j) = Cov\left[N_{ij}^{(s)}, N_{ij}^{(s)}\right] = Var\left(N_{ij}^{(s)}\right)$. Because $N_{ij}^{(s)}$ follows a Poisson distribution, $Var\left(N_{ij}^{(s)}\right) = \lambda_{ij}^{(s)}$, and:

$$Cov\left(N_i, N_j\right) = vLT_{ij}^{(s)} \qquad \text{Equation (1)}$$

##### 4.3. Covariance of the number of mutations carried by two diplotypes

Let $\{ij\}$ and $\{hk\}$ denote two diplotypes formed by the pairs of haplotypes $i$, $j$ and $h$, $k$, respectively. The number of mutations carried by them is given by $N_{\{i\ j\}} = N_i + N_j$, and $N_{\{hk\}} = N_h + N_k$. Then the covariance of the number of mutations carried by them is:

$$\begin{aligned} Cov\left(N_{\{ij\}}, N_{\{jk\}}\right) &= Cov\left[\left(N_i + N_j\right), \left(N_h + N_k\right)\right] \\ &= Cov(N_i, N_h) + Cov(N_i, N_k) + Cov\left(N_j, N_h\right) + Cov\left(N_j, N_k\right) \end{aligned}$$

Using equation (1), this expression reduces to:

$$Cov(N_{\{i,j\}}, N_{h,k\}}) = vLT_{ih}^{(s)} + vLT_{ik}^{(s)} + vLT_{jh}^{(s)} + vLT_{jk}^{(s)} \qquad \text{Equation (2)}$$

##### 4.4. Correlation of the number of mutations carried by two diploid individuals

By definition:

$$Cor\left(N_{\{ij\}}, N_{\{hk\}}\right) = \frac{Cov\left(N_{\{ih\}}, N_{\{hk\}}\right)}{\sqrt{Var\left(N_{\{ij\}}\right) \times Var\left(N_{\{hk\}}\right)}} \qquad \text{Equation (3)}$$

To obtain the denominator, we note that $N_{\{ij\}} = N_i + N_j$, and $N_{\{hk\}} = N_h + N_k$. Note that $T_{ii}^{(s)} = T^{(R)}$ for any haplotype $i$. Thus:

$$\begin{aligned} Var\left(N_{\{i,j\}}\right) &= Var(N_i) + Var\left(N_j\right) + 2Cov\left(N_i, N_j\right) \\ &= 2vL\left(T^{(R)} + T_{ij}^{(s)}\right) \\ &= 2vL\left(T_R + T_{hk}^{(s)}\right) \end{aligned} \qquad \text{Equation (4)}$$

And analogously:

$$Var\left(N_{\{h,k\}}\right) = 2vL\left(T_R + T_{hk}^{(s)}\right)$$

We replace Equations (2) and (4**Error! Reference source not found.**) into Equation (3) and obtain:

$$\begin{aligned} GeSi_{Tree} &:= Cor(N_{\{i,j\}}, N_{\{h,k\}}) \\ &= \frac{vL\left(T_{ih}^{(s)} + T_{ik}^{(s)} + T_{jh}^{(s)} + T_{jk}^{(s)}\right)}{2vL\sqrt{\left(T^{(R)} + T_{ij}^{(s)}\right)\left(T^{(R)} + T_{hk}^{(s)}\right)}} \\ &= \frac{T_{ih}^{(s)} + T_{ik}^{(s)} + T_{jh}^{(s)} + T_{jk}^{(s)}}{2\sqrt{\left(T^{(R)} + T_{ij}^{(s)}\right)\left(T^{(R)} + T_{hk}^{(s)}\right)}} \end{aligned} \qquad \text{Equation (5)}$$

A disadvantage of $GeSi_{Tree}$ is that it requires knowing the true genealogy. We will later demonstrate how to calculate it from genotype data without inferring the genealogical tree.

#### 5. $GeSi_{Tree}$ as the correlation of the deviations from the ancestral phenotype due to additive genetic effects

We now demonstrate that $GeSi_{Tree}$ also represents the correlation of the deviation of genetics effects from that of a hypothetical ancestor. To model the deviation from the ancestral phenotype due to total genetic effects, we need to focus on the genetic variants with a causal effect on a trait. Thus, the parameters $N_i$, $N_j$, $N_h$, and $N_k$ now represent the number of causal variants carried by the respective haplotypes, and the $\lambda$ parameters denote the rate at which causal mutations arise. We will need to assume that the number of causal mutations is linear with time. For now, we will also assume that the effect sizes of all variants are sampled from the same distribution and are independent of the minor allele frequency, similar to the LDAK model with $\alpha = 0$.

##### 5.1. Variance of the phenotypic deviation

First, we express the phenotypes as deviations from the phenotype of the ancestral haplotype. Let $Y_{\{i,j\}}$ and $Y_{\{h,k\}}$ denote the phenotypes of two individuals carrying the diplotypes $\{i,j\}$ and $\{h,k\}$, respectively. We also define $d_i$ as a deviation from the ancestral phenotypic value caused by any haplotype $i$. Then, the phenotypes can be expressed as:

$$Y_{\{ij\}} = y_0 + d_i + d_j + e_{\{ij\}}$$

$$Y_{\{hk\}} = y_0 + d_h + d_k + e_{\{hk\}}$$

Where $y_0$ is the phenotypic value of a hypothetical individual carrying two copies of the ancestral haplotype. We want to model the correlation between $Y_{\{i,j\}}$ and $Y_{\{h,k\}}$ in terms of the divergence. We assume and additive model and define $d_i$, for any haplotype $i$, as the sum of the causal effect sizes "a":

$$d_i = \sum_{m=1}^{N_i} a_m \qquad \text{Equation (6)}$$

We assume the effect sizes follow some distribution with mean $\mu_a$, and variance $\sigma_a^2$. We apply the law of total variance to find the variance of *d* in equation (6**Error! Reference source not found.**):

$$Var(d_i) = E[Var(d_i|N_i)] + Var(E[d_i|N_i])$$

We simplify the first term on the right-hand side by assuming that the effect sizes of two variants on a haplotype are uncorrelated:

$$\begin{aligned} Var(d_i) &= E[\sigma_a^2 N_i] + Var(\mu_a N_i) \\ &= \sigma_a^2 \lambda_i + \mu_a^2 \lambda_i = \lambda_i(\sigma_a^2 + \mu_a^2) \\ &= vL(\sigma_a^2 + \mu_a^2)T^{(R)} \end{aligned}$$

This expression holds true for any time interval during which the number of mutations is linear with time. Thus, in general:

$$Var(d) = vL(\sigma_a^2 + \mu_a^2)T \qquad \text{Equation (7)}$$

Where $T$ is any arbitrary time interval where this assumption holds true, and $d$ is the phenotypic deviation caused by the mutations that occurred during that time.

##### 5.2. Covariance of the phenotypic deviation terms

We now want to model the covariance between the phenotypic divergence caused by two haplotypes $i$ and $j$. We use the same approach as in Section 4.2 and express each deviation term as the sum of shared and unshared deviations between any pair of haplotypes $i$ and $j$:

$$d_i = d_{ij}^{(s)} + d_i^{(u)} = \sum_{m=1}^{N_{ij}^{(s)}} a_m + \sum_{q=1}^{N_i^{(u)}} a_q$$

$$d_j = d_{ij}^{(s)} + d_j^{(u)} = \sum_{m=1}^{N_{ij}^{(s)}} a_m + \sum_{q=1}^{N_j^{(u)}} a_q$$

Where the $d_{ij}^{(s)}$ represents the shared phenotypic divergence due shared mutations $N_{ij}^{(s)}$ between haplotypes $i$ and $j$, and $d_i^{(u)}$ and $d_j^{(u)}$ represent the unshared phenotypic divergences of haplotypes $i$ and $j$ due to unshared mutations $N_i^{(u)}$ and $N_j^{(u)}$, respectively. The covariance between $d_i$ and $d_j$ is given by:

$$Cov(d_i, d_j) = Cov\left[\left(\sum_{m=1}^{N_{ij}^{(s)}} a_m + \sum_{q=1}^{N_i^{(u)}} a_q\right), \left(\sum_{m=1}^{N_{ij}^{(s)}} a_m + \sum_{q=1}^{N_j^{(u)}} a_q\right)\right]$$

$$= Var\left(\sum_{m=1}^{N_{ij}^{(s)}} a_m\right) + Cov\left[\left(\sum_{m=1}^{N_{ij}^{(s)}} a_m\right), \left(\sum_{q=1}^{N_j^{(u)}} a_q\right)\right] +$$

$$Cov\left[\left(\sum_{q=1}^{N_i^{(u)}} a_q\right), \left(\sum_{m=1}^{N_{ij}^{(s)}} a_m\right)\right] + Cov\left[\left(\sum_{q=1}^{N_i^{(u)}} a_q\right), \left(\sum_{q=1}^{N_j^{(u)}} a_q\right)\right]$$

$$= Var\left(\sum_{m=1}^{N_{ij}^{(s)}} a_m\right)$$

$$= Var\left(d_{i,j}^{(s)}\right) \qquad \text{Equation (8)}$$

The covariance terms were cancelled under the assumption that the phenotypic deviation of two haplotypes from the ancestral phenotype is independent of each other once their genealogies have diverged. By replacing Equation (7) into Equation (8) and using the appropriate time parameter, we obtain:

$$Cov(d_i, d_j) = vL(\sigma_a^2 + \mu_a^2)T_{i,j}^{(s)} \qquad \text{Equation (9)}$$

Thus, the covariance of phenotypic deviations is equal to the variance of the shared deviation.

#### 5.3. Correlation of the phenotype of two diploid individuals

To calculate the correlation, we need to calculate the variance and the covariance. We start by modeling the variance of the phenotype of a diploid individual. As per Section 4.1:

$$d_{\{ij\}} = y_0 + d_i + d_j + e_{\{i,j\}}$$

Then:

$$\begin{aligned} Var(d_{\{ij\}}) &= Var(d_i + d_j) \\ &= Var(d_i) + Var(d_j) + 2Cov(d_i, d_j) \end{aligned}$$

By plugging in the results from Equation (7) and Equation (9), we obtain:

$$\begin{aligned} Var(d_{\{ij\}}) &= 2vLT^{(R)}\,(\sigma_a^2 + \mu_a^2) + 2vLT_{ij}^{(s)}\,(\sigma_a^2 + \mu_a^2) \\ &= 2vL\,(\sigma_a^2 + \mu_a^2)\,(T^{(R)} + T_{ij}^{(s)}) \end{aligned} \qquad \text{Equation (10)}$$

The covariance of $d_{\{i,j\}}$ and $d_{\{h,k\}}$ is given by:

$$\begin{aligned} Cov(d_{\{ij\}}, d_{\{hk\}}) &= Cov(y_0 + d_i + d_j + e_{\{ij\}}, y_0 + d_h + d_k + e_{\{h,k\}}) \\ &= Cov(d_i, d_h) + Cov(d_i, d_k) + Cov(d_j, d_h) + Cov(d_j, d_k) \end{aligned}$$

$$= vL(\sigma_a^2 + \mu_a^2)\left(T_{ih}^{(s)} + T_{ik}^{(s)} + T_{jh}^{(s)} + T_{jk}^{(s)}\right) \qquad \text{Equation (11)}$$

And the variance is given by:

$$\begin{aligned} Var\left(d_{\{i,j\}}\right) &= Cov\left(d_{\{i,j\}}, d_{\{i,j\}}\right) \\ &= vL(\sigma_a^2 + \mu_a^2)\left(T_{ii}^{(s)} + T_{ij}^{(s)} + T_{ij}^{(s)} + T_{jj}^{(s)}\right) \\ &= 2vL(\sigma_a^2 + \mu_a^2)\left(T^{(R)} + T_{ij}^{(s)}\right) \qquad \text{Equation (12)} \end{aligned}$$

Finally, to obtain the correlation, we standardize the covariance by the square root of the product of the variances:

$$\begin{aligned} Cor(d_{\{ih\}}, d_{\{hk\}}) &= \frac{Cov\left(d_{\{ij\}}, d_{\{hk\}}\right)}{\sqrt{Var(d_{\{ij\}})Var(d_{\{hk\}})}} \\ &= \frac{vL(\sigma_a^2 + \mu_a^2)\left(T_{ih}^{(s)} + T_{ik}^{(s)} + T_{jh}^{(s)} + T_{jk}^{(s)}\right)}{\sqrt{\left(2vL\,(\sigma_a^2 + \mu_a^2)\,\left(T^{(R)} + T_{ij}^{(s)}\right)\right)\left(2vL\,(\sigma_a^2 + \mu_a^2)\left(T^{(R)} + T_{hk}^{(s)}\right)\right)}} \\ &= \frac{vL(\sigma_a^2 + \mu_a^2)\left(T_{ih}^{(s)} + T_{ik}^{(s)} + T_{jh}^{(s)} + T_{jk}^{(s)}\right)}{2vL\,(\sigma_a^2 + \mu_a^2)\,\sqrt{\left(T^{(R)} + T_{ij}^{(s)}\right)\left(T^{(R)} + T_{hk}^{(s)}\right)}} \\ &= \frac{\left(T_{ih}^{(s)} + T_{ik}^{(s)} + T_{jh}^{(s)} + T_{jk}^{(s)}\right)}{2\sqrt{\left(T^{(R)} + T_{ij}^{(s)}\right)\left(T^{(R)} + T_{hk}^{(s)}\right)}} \end{aligned}$$

The expression above is the same as the definition of $GeSi_{Tree}$ given in Equation (**Error! Reference source not found.**5). Thus, $GeSi_{Tree}$ also represents the correlation of the deviation from the expected ancestral phenotype under the assumption that LDAK's $\alpha = 0$, and the number of causal mutations carried by a haplotype is linear with time:

$$\begin{aligned} Cor(d_{\{i,h\}}, d_{\{h,k\}}) &\equiv GeSi_{Tree} \\ &= \frac{T_{ih}^{(s)} + T_{ik}^{(s)} + T_{jh}^{(s)} + T_{jk}^{(s)}}{2\sqrt{\left(T^{(R)} + T_{ij}^{(s)}\right)\left(T^{(R)} + T_{hk}^{(s)}\right)}} \qquad \text{Equation (13)} \end{aligned}$$

##### 5.4. Calculating $GeSi_{tree}$ from genotype data

We now demonstrate that GeSi can be calculated directly from genotype data without inferring the genealogy. First, we note that the derivation of Equations (5) and (13) relied on a Poisson approximation of the number of shared mutations given the shared divergence time. We also note that each of the right-hand terms of Equation (11) represented the lambda parameter of such Poisson distributions:

$$Cov(N_{\{i,j\}}, N_{\{h,k\}}) = vLT_{ih}^{(s)} + vLT_{ik}^{(s)} + vLT_{jh}^{(s)} + vLT_{jk}^{(s)}$$

$$= \lambda_{ih}^{(s)} + \lambda_{ik}^{(s)} + \lambda_{jh}^{(s)} + \lambda_{jk}^{(s)}$$

For a Poisson process, the observed mutation counts are both the maximum likelihood and the method-of-moments estimators of the lambda parameters. Thus, we replace in the realized number of mutations and obtain:

$$\widehat{Cov}(N_{\{ij\}}, N_{\{hk\}}) \quad = N_{ih}^{*(s)} + N_{ik}^{*(s)} + N_{jh}^{*(s)} + N_{jk}^{*(s)} \qquad \text{Equation (14)}$$

Where the $N^{*(s)}$ parameters have the same interpretation as the $N^{(S)}$ parameters before, except that they represent the realized counts of derived alleles rather than random variables.

The advantage of Equation (14) is that it can be calculated from whole genome sequence data without knowing the tree, provided the reference allele is set to the ancestral allele, and the alternate allele to the derived allele, and the analysis is limited to biallelic sites. To see this, we first express each of the $N^{*(S)}$ parameters as a sum of indicator variables:

$$N_{ih}^{*(s)} = \sum_{m=1}^{p} I_{ih,m}^{*}$$

Where $m$ is the position along a sequence of length $p$, and $I^*$ is an indicator variable that takes a value of 1 if both haplotypes $i$ and $h$ carry the derived allele at position $m$ and takes the value 0 otherwise. Importantly, this expression holds true even under linkage disequilibrium (LD) because of the linearity of the summation operator. We apply the same expression to the other terms in Equation (13) and obtain:

$$\begin{aligned} \widehat{Cov}(N_{\{ij\}}, N_{\{hk\}}) \quad &= \sum_{m=1}^{p} I_{ih,m}^{*} + \sum_{m=1}^{p} I_{ik,m}^{*} + \sum_{m=1}^{p} I_{jh,m}^{*} + \sum_{m=1}^{p} I_{jk,m}^{*} \\ &= \sum_{m=1}^{p} S_m^{*} \qquad \text{Equation (15)} \end{aligned}$$

Where:

$$S_m^{*} \quad := \; I_{ih,m}^{*} + I_{ik,m}^{*} + I_{jh,m}^{*} + I_{jk,m}^{*} \qquad \text{Equation (16)}$$

$S_m^*$ in Equation (16) is the number of shared derived alleles at a single site $m$. To facilitate calculation directly from genotype data, we introduce matrix notation indexing by subject and site. Let $\mathbf{X} \in \{0,1,2\}^{n \times p}$ denote the genotype matrix for $n$ subjects across $p$ sites, where entries $x_{i,m}$ represent the number of derived alleles for subject $i$ at site $m$. Let subjects $a, b \in \{1,2,\ldots,n\}$ carry the diplotypes $\{ij\}$ and $\{hk\}$, respectively. The realized total number of derived alleles carried by them is given by $N_{\{ij\}}^{*} = \sum_{m=1}^{p} x_{am}$, and $N_{\{hk\}}^{*} = \sum_{m=1}^{p} x_{bm}$. In Suppl. Table 1, we tabulate both the pairwise number of shared derived alleles and the product of the coded genotypes and demonstrate that the left-hand side of equation (16) is numerically equivalent to $x_{am}x_{bm}$. We substitute this identity on Equation (15) and obtain:

$$\widehat{Cov}(N_{\{ij\}}, N_{\{hk\}}) = \sum_{m=1}^{p} S_m^* = \sum_{m=1}^{p} x_{am}x_{bm} = \mathbf{x}_a \cdot \mathbf{x}_b \qquad \text{Equation (17)}$$

Where $\mathbf{x}_a, \mathbf{x}_b \in \{0,1,2\}^p$ are the genotype vectors for subjects $a$ and $b$.

In Equation (5), we defined $GeSi_{tree}$ as the correlation of the number of derived alleles between two diploid individuals, and in equation (13) we demonstrated that $\mathrm{GeSi}_{tree}$ also represents their phenotype correlation under specific assumptions (LDAK's $\alpha = 0$, and the number of causal sites is linear with time). We now aim to estimate the $n \times n$ matrix $\widehat{\mathbf{GeSi}}_{\mathrm{tree}}$ of estimated $GeSi_{tree}$ coefficients directly from the genotype matrix $\mathbf{X}$. The strategy is to first obtain the variance-covariance matrix $\hat{\mathbf{C}}$, whose entry $(ab)$ estimates the covariance term in the numerator of Equation (13**Error! Reference source not found.**). We then use the diagonal elements of $\hat{\mathbf{C}}$, which estimate the variance terms in the denominator of Equation (13), to standardize the matrix. Equation (17) established that the $(a, b)$-th entry of the covariance matrix $\hat{\mathbf{C}}$ is $\mathbf{x}_a \cdot \mathbf{x}_b$. Therefore, the full variance-covariance matrix is:

$$\hat{\mathbf{C}} = \mathbf{X}\mathbf{X}^T$$

Let $\mathbf{D} = \mathrm{diag}(\hat{\mathbf{C}})$ be diagonal matrix of variances. The $\widehat{\mathbf{GeSi}}_{\mathrm{tree}}$ matrix is then obtained by standardizing $\hat{\mathbf{C}}$:

$$\widehat{\mathbf{GeSi}}_{\mathrm{tree}} = \mathbf{D}^{-1/2}\,\hat{\mathbf{C}}\mathbf{D}^{-1/2} = \mathbf{D}^{-1/2}\mathbf{X}\mathbf{X}^T\mathbf{D}^{-1/2} \qquad \text{Equation (18)}$$

**Supplementary table 1.** Estimation of Equation (15) using genotype data

| Subject 1 | | | Subject 2 | | | Estimation of Equation (15) | |
|---|---|---|---|---|---|---|---|
| Status | Genotype | $g_{m\{i,j\}}$ | Status | Genotype | $g_{m\{h,k\}}$ | $S_m^*$ | $x_{a,m} \times x_{b,m}$ |
| Hom. Anc. | Anc/Anc | 0 | Hom. Anc. | Anc/Anc | 0 | 0 | 0 |
| Hom. Anc. | Anc/Anc | 0 | Het. | Anc/Der | 1 | 0 | 0 |
| Hom. Anc. | Anc/Anc | 0 | Hom. Der. | Der/Der | 2 | 0 | 0 |
| Het. | Anc/Der | 1 | Het. | Anc/Der | 1 | 1 | 1 |
| Het. | Anc/Der | 1 | Hom. Der. | Der/Der | 2 | 2 | 2 |
| Hom. derived | Der/Der | 2 | Hom. Der. | Der/Der | 2 | 4 | 4 |

Hom: Homozygous. Het: Heterozygous. Anc: Ancestral allele. Der: Derived allele. $S_m$ defined in Equation (8).

##### 5.5. Geometric interpretation and extension of the GeSi model

The estimator $\widehat{GeSi}_{tree}$ in Equation (18) represents a matrix of cosine similarities between the raw genotype vectors:

$$\widehat{GeSi}_{tree(ab)} \equiv \frac{\mathbf{x}_a . \mathbf{x}_b}{|\mathbf{x}_a||\mathbf{x}_b|} \qquad \text{Equation (19)}$$

Thus, $\widehat{GeSi}_{tree(ab)}$ measures the cosine of the angle between the raw genotype vectors originating from the zero vector (i.e. the root's genotype vector). Thus, we conclude that the genotype cosine similarity is an estimator of the phenotype Pearson correlation due to genetic effects. Note that this differs from standard GRM methods, which use a weighted average of per-site genotype covariances and thus require LD-pruned genotype data. Thus, within the genomic region spanned by a single genealogical tree, calculating the GeSi coefficient does not require LD pruning, provided the derived sites are uniformly distributed along the sequence. We note however that real genomic data is usually spanned by more than one genealogical tree and that the assumption of uniformly distributed sites is likely violated, and thus LD-pruning or thinning might be required, hence why all our benchmarks included WGS, LD-pruned and randomly selected sites data.

The original derivation of GeSi, $GeSi_{tree}$, and its estimator $\widehat{\mathbf{GeSi}}_{\text{tree}}$, modeled the Pearson correlation of the deviations from the ancestor's total genetic effects ($d_{\{ij\}} = g_a$) due to the total divergence time from the present to the root of the tree. Importantly, it relied on a Poisson distribution to model the variance that would be created if we followed all possible lineages during a given amount of time. However, study samples are not representative of all potential lineages and thus a sample variance will not match the variance parameters used by $GeSi_{tree}$.

To address this limitation and focus on the variance observed in an ascertained sample, we generalize the geometric interpretation. We hypothesize that the genotype cosine similarity can model the Pearson correlation of genetic effects relative to reference points other than the ancestral origin. A natural reference for an ascertained sample is its centroid, defined by the mean genotype vector $\bar{\mathbf{x}}$. Importantly, this model implies a potential mechanism of why heritability estimates are not transferable between populations.

Let $n$ denote the number of subjects in a very large set of samples representative of all potential lineages descending from the root of a tree. Let $S = \{1, \ldots, n_S\}$ with $n_S < n$ denote the set of all ascertained subjects included in a study. Let $\mathbf{X} \in \mathbb{R}^{n \times p}$ denote the raw genotype matrix of a very large sample representative of all potential lineages deriving from the root of a tree. Let $\mathbf{X}_S \in \mathbb{R}^{n_S \times p}$ denote the genotype matrix of the ascertained subjects. Let $\bar{\mathbf{x}}_{asc} \in \mathbb{R}^p$ denote the vector of mean genotypes in the ascertained set. Let $\mathbf{W} = \mathbf{X} - \mathbf{1}_n \bar{\mathbf{x}}_{asc}^T \in \mathbb{R}^{n \times p}$ be the full genotype matrix centered on the ascertained samples' mean genotype, and $\mathbf{W}_S \in \mathbb{R}^{n_S \times p}$ a subset of $\mathbf{W}$ containing only the ascertained subjects. Let $\mathbf{w}_a$ and $\mathbf{w}_b$, where $a, b \in \{1, \ldots, n\}$, denote the $a$-th and $b$-th row of $\mathbf{W}$, respectively. Then the angle, and therefore the cosine similarity, between the displacement vectors $(\mathbf{x}_a - \bar{\mathbf{x}}_{asc})$ and $(\mathbf{x}_b - \bar{\mathbf{x}}_{asc})$ originating from the ascertained samples' centroid $\bar{\mathbf{x}}_{asc}$ in the original space $\mathbf{X}$ are identical to the angle and cosine similarity between the vectors $\mathbf{w}_a$ and $\mathbf{w}_b$ originating from the origin in the centered space. Notably, $\widehat{\mathbf{GeSi}}_{tree} = \mathbf{D}^{-1/2}\mathbf{X}\mathbf{X}^T\mathbf{D}^{-1/2}$, which represents the cosine similarity in the original space $\mathbf{X}$, is an estimator of the Pearson correlation of the total genetic effects measured as deviations from the ancestral phenotype at the root of the tree. Therefore, we hypothesize that $\widehat{\mathbf{GeSi}}_{asc}$, the cosine similarity between the centered vectors in the $\mathbf{W}$ space, are estimators of the Pearson correlation of genetic effects of the ascertained samples:

$$\widehat{\mathbf{GeSi}}_{asc} = \mathbf{D}_{\mathbf{W}}^{-1/2}\mathbf{W}\mathbf{W}^T\mathbf{D}_{\mathbf{W}}^{-1/2}$$

Where $\mathbf{D}_{\mathbf{W}} = \text{diag}(\mathbf{W}\mathbf{W}^T)$. The GeSi coefficient for any two samples depends only on those samples' genotype vectors. Thus, we can calculate GeSi for just the ascertained samples by restricting the genotype matrix to contain only the ascertained samples:

$$\widehat{\mathbf{GeSi}}_{asc,S} = \mathbf{D}_{\mathbf{W}_S}^{-1/2}\mathbf{W}_S\mathbf{W}_S^T\mathbf{D}_{\mathbf{W}_S}^{-1/2}$$

Where $\widehat{\mathbf{GeSi}}_{asc,S} \in \mathbb{R}^{n_S \times n_S}$ is an estimator of the genetics effects deviations from the ascertained samples' mean. The inherent dependence on the sample mean implies that the variance components are sample-specific and would contribute to explaining the observed lack of transferability of heritability components across populations.

We further extend GeSi to allow for differential contributions of different sites based on, e.g., minor allele frequency, such as is the case for LDAK. Let $f_m$ denote the scaling factor for site $m$, where $m \in \{1, \dots, p\}$. To obtain a LDAK-like scaling, we set $q_m = [2f_m(1 - f_m)]^{\alpha/2}$, where $f_m$ is the alternate allele frequency at site $m$ in the ascertained sample, and $\alpha$ an arbitrary scalar. Let $\mathbf{Q} = \mathrm{diag}(q_1, \dots, q_p)$ be a diagonal matrix containing the scaling factors. Then the matrix of scaled genotypes is $\mathbf{Z_S} = \mathbf{W_S Q}$. We hypothesize that if these scaling factors reflect the true causal relationship between genotype and phenotype, then the cosine similarity calculated in this scaled space is an estimator of the Pearson correlation of the genetic effects deviations from the ascertained samples' mean.

$$\widehat{\mathbf{GeSi}}_{asc,S,\alpha} = \mathbf{D}_{\mathbf{Z}_S}^{-1/2} \mathbf{Z}_S \mathbf{Z}_S^T \mathbf{D}_{\mathbf{Z}_S}^{-1/2} \qquad \text{Equation (20)}$$

Because in practice we only have access to the ascertained sample, for the rest of this manuscript we drop the subscripts related to the ascertainment process and refer to the estimator in Equation (20) as $\widehat{\mathbf{GeSi}}_{\alpha}$.

##### 5.6. Using GeSi to estimate variance components via mixed linear models

Standard GRM methods assume that a single genetic variance component exists and is estimable through mixed linear models (MLMs). However, the generative model in Equation (13) showed that the variance of a subject's genetic effects depends on its amount of autozygosity. If this model held true, it would mean that MLMs cannot estimate a single genetic variance parameter because such parameter does not exist. Instead, MLMs would estimate an average of the pairwise product of genetic effects standard deviations:

$$Cor(g_i, g_j) = GeSi_{tree}$$

$$\frac{Cov(g_i, g_j)}{\sigma_{g_i}\sigma_{g_j}} = GeSi_{tree}$$

$$Cov(g_i, g_j) = GeSi_{tree} \times \left[\sigma_{g_i}\sigma_{g_j}\right]$$

Thus, to use $\widehat{\mathbf{GeSi}}_{\alpha}$ in a mixed linear model, we need to assume that a single genetic variance component exists, in which case:

$$\widehat{Cov}(g_i', g_j') = \widehat{\mathrm{GeSi}}_{\alpha} \times \sigma_g^2$$

##### 5.7. Calculation of GeSi in the real and simulated genotype data

$GeSi_{tree}$ was defined based on a single genealogical tree, however, the genealogy of DNA sequences that have undergone recombination cannot be represented by a single tree. Thus, we calculated the genome-wide $\mathbf{GeSi}_{tree}$ value as the average of all trees along the genome weighted by the tree span, i.e. the length of the genomic region covered by a tree. $\widehat{\mathbf{GeSi}}_{tree}$, the genotype-based estimator of $\mathbf{GeSi}_{tree}$, was calculated using all biallelic SNPs in whole-genome sequence (WGS) data, or using only sites that were roughly in linkage equilibrium (LD-pruned), or using sites selected at random with uniform probability (RS). We assessed the concordance between $\mathbf{GeSi}_{tree}$ and $\widehat{\mathbf{GeSi}}_{tree}$ by calculating the coefficient of determination ($R^2$) and the Pearson's correlation coefficient, by plotting them against each other, and by visually comparing heatmaps of the distribution of GeSi values calculated by the two approaches. After validating $\widehat{\mathbf{GeSi}}_{tree,WGS}$ as an estimator of $\mathbf{GeSi}_{tree}$, we visualized the distribution of $\widehat{\mathbf{GeSi}}_{tree,WGS}$ values of related and unrelated pairs of subjects through violin plots stratified by ancestry pairs.

Finally, we calculated $\widehat{\mathbf{GeSi}}_{\alpha}$ using WGS, LD-pruned or RS genotype data with values of alpha ranging between 0 and -2 with steps of -0.25.

The GeSi implementation and documentation is available at https://github.com/diegovelizo/gesi.

#### 6. Estimation of genetic similarity matrices (GRMs)

##### 6.1. LDAK and GCTA GRMs

We generated Genetic Relationship Matrices (GRMs) using LDAK[11] v.6.1 with parameter $\alpha$ values ranging from -2 to 0 in increments of -0.25. The GRMs with $\alpha = -1$ are equivalent to the GCTA GRM. For each dataset, we used up to three different data types: All biallelic SNPs in whole-genome sequence data ("WGS"), a subset of sites with low levels of linkage disequilibrium ("LD-pruned"), and randomly selected sites ("RS"). The LD-pruned subset was generated using PLINK[17] v.1.9 with the option --indep-pairwise. For the Simulation Sets #2 and #3 (Msprime-generated), the window size was set to 400 SNPs with steps of 50 SNPs, and r2 threshold of 0.1. For the BioMe data, the window size was 1000 SNPs with step of 100 SNPs, and r2 threshold of 0.07. RS SNPs were selected with uniform probability from the biallelic SNPs WGS data, ensuring the number of sites per chromosome was proportional to the total WGS SNP count per chromosome. The total number of RS sites across all autosomes was the same as the total number of sites in the corresponding LD-pruned set.

For the LD-pruned and RS datasets, a single GRM corresponding to all autosomes was computed for each $\alpha$. For the full WGS data, GRMs were first calculated independently for each autosome and subsequently averaged in R, weighting each chromosome-specific GRM by its respective SNP count, to generate the final genome-wide GRM for each $\alpha$.

For the LD-pruned and RS datasets, PCs were calculated directly with LDAK via the --pca option keeping 20 PCs. For the WGS data, PCs were obtained via Eigen decomposition (eigen function in R) of each final averaged GRM (one GRM per $\alpha$ value per data type per dataset). For each decomposition, the top 20 eigenvectors were retained as the PCs.

##### 6.2. PC-Relate and KING-robust GRMs

Estimating the PC-Relate[18] GRM required a series of iterative steps. First, we calculated an initial GRM using the KING-robust method[19] via SNPRelate::snpgdsIBDKING (i.e. the snpgdsIBDKING function in the R package SNPRelate). Next, we calculated the principal components via the PCAiR method implemented in GENESIS::pcair using the KING-robust matrix as input, with a kinship threshold of $(1/2)^{4.5}$ and divergence threshold of $-(1/2)^{4.5}$, corresponding to the maximum kinship coefficient value for third-degree relatives[9]. Next, we calculated the PC-Relate kinship matrix via the GENESIS::pcrelate function, using the first four principal components calculated with PCAiR in the previous step as covariates. We ran a second and final round of PCAiR using the PC-Relate kinship estimates as input and keeping the kinship threshold at $(1/2)^{4.5}$ and the divergence threshold at $-(1/2)^{4.5}$. The output of PCAiR included a list of unrelated and related individuals based on the PC-Relate kinship estimates and the specified kinship threshold.

#### 6.3. Mixed linear model association testing

We used mixed linear models (MLMs) for both genome-wide association studies (GWAS) and phenotype prediction. The general model was $\mathbf{Y} = \mathbf{X\beta} + \mathbf{Zu} + \mathbf{e}$, where $\mathbf{Y}$ is the phenotype vector of length $n$, $\mathbf{X} \in \mathbb{R}^{n \times p}$ is the design matrix for fixed effects, $\boldsymbol{\beta}$ is the vector of fixed effect sizes of length $p$, $\mathbf{Z}$ is the design matrix for random effects, $\mathbf{u}$ is the vector of random genetic effects, and $\mathbf{e}$ is the residual error vector. We assumed that $e \sim N(0, \mathbf{I}\sigma_e^2)$ and $\mathbf{u} \sim N(\mathbf{0}, \sigma_a^2\, \mathbf{K})$, where $\mathbf{K}$ is a genetic similarity matrix (either a GRM or a GeSi matrix) and $\sigma_a^2$ is the additive genetic variance, and $\sigma_e^2$ is the residual variance.

We ran a GWAS of the using the GENESIS package for R in a two-step procedure. In the first step, we fit the null model through the function `GENESIS::fitNullModel`, with all the fixed-effect covariates and random effect matrices but excluding single genetic variants. In the second step, we ran the single-variant association tests using the null model from the previous step through the function `GENESIS::assocTestSingle`. We used the Gaussian family for quantitative traits, and the binomial family for binary traits. We used the saddlepoint approximation to calibrate the score test statistic for binary traits [20,21].

All models were run with either ten or zero Principal Components (PCs) as fixed effects. GWAS of Simulated Sets #2 and #3 did not include any covariates other than PCs. GWAS of Simulated Sets #4-7 and the real phenotype data from the BioMe cohort included sex, age, age$^2$, population label, as covariates. The residuals were assumed to be homoscedastic in all GWAS of Simulated Sets #2-7.

GWAS of height and BMI data from the BioMe cohort used a fully-adjusted two-stage residual rank normalization[22] (options `two.stage=TRUE` and `norm.option="by.group"` in the `GENESIS::assocTestSingle` function), and were run either assuming that the residuals were homoscedastic, or allowing for six different residual variance components, one per each sex x population label interaction (option `group.var="ancestry_sex"`).

The heritability was estimated in all datasets through the function `GENESIS::fitNullModel`. The single-variant association tests were run only in Simulated Sets #2-3. We used the Simulated Sets #2-3 to systematically assess:

1. **Type I error rate**: The proportion of truly null SNPs declared significant under a nominal p-value threshold ($p < 0.05$) or genome-wide significance ($p < 5\times10^{-8}$).
2. **Statistical power**: The proportion of truly causal SNPs achieving significance, particularly relevant when comparing how different methods handle admixture or cryptic relatedness.
3. **Genomic inflation**: Calculated as the genomic control lambda ($\lambda_{GC}$) statistic.
4. **Effect size accuracy**: Measured the mean squared error of the estimates effect sizes, and by the squared Pearson correlation ($r^2$) and the coefficient of determination ($R^2$) between true and estimated effect sizes.
5. **Effect size bias**: Assessed by the regression coefficient of the estimated effect sizes on the true effect sizes.
6. **Heritability estimation**: Mean squared error of the heritability estimates, by comparing the inferred values against the simulated parameters.

##### 6.4. Inference of the $\alpha$ parameter through CV-BLUP

We used the Best Linear Unbiased Predictor (BLUP) equation[23,24] to predict the total genetic effects in the held-out sets of a 20-fold cross-validation (CV) analysis of Simulated Sets #4-7 and the height and BMI traits from BioMe. We repeated this analysis for each value of the $\alpha$ parameter values, in order to assess whether this approach could be used to find its true value. Additionally, we recorded the restricted maximum likelihood of all models as an alternative approach to select the best $\alpha$ value, as has been done before [25].

All datasets were split into 20 cross-validation folds of 494 subjects each. Because the total sample size (9,885) was not a multiple of 20, we randomly excluded the same five samples from all cross-validation analyses. We fit a MLM the `GENESIS::fitNullModel` function on the training set of each cross-validation fold. The null models included a genetic similarity matrix (either a GRM or a GeSi matrix) and the same covariates as in the MLM association testing. Thus, we obtained the variance components estimates for each fold, which were then fed into the BLUP equation:

$$\hat{\mathbf{u}} = \mathbf{G}\mathbf{Z}^{\mathrm{T}}\mathbf{V}^{-1}(\mathbf{Y} - \mathbf{X}\hat{\boldsymbol{\beta}})$$

Where $\mathbf{Y} \in \mathbb{R}^{n\times 1}$ is the vector of phenotypes of the training set (n = 9,386); $\mathbf{X} \in \mathbb{R}^{n\times p}$ is the matrix of fixed effects covariates in the training set $\hat{\boldsymbol{\beta}} \in \mathbb{R}^{p\times 1}$ is the vector of fixed effects coefficients estimated in the training set; $\hat{\mathbf{u}} \in \mathbb{R}^{(n+m)\times 1}$ is the predicted genetic effects of both the training and validation (m = 494) sets; $\mathbf{G} = \sigma_a^2\mathbf{K}$ is the estimated covariance matrix of the genetic effects; $\mathbf{K}$ is the genetic similarity matrix of both training and validation sets, and $\mathbf{G}, \mathbf{K} \in \mathbb{R}^{(n+m)\times(n+m)}$. The incidence matrix $\mathbf{Z} \in \mathbb{R}^{n\times(n+m)}$ maps the genetic effects to each subject, $\mathbf{V} = (\mathbf{Z}\mathbf{G}\mathbf{Z}^{\mathrm{T}} + \mathbf{R})$, where $\mathbf{V} \in \mathbb{R}^{n\times n}$ is the total variance matrix, and $\mathbf{R} \in \mathbb{R}^{n\times n}$ is the diagonal matrix of residual variance.

We calculated the total phenotype of the validation set as the sum of estimated fixed effects and random genetic effects $\hat{\mathbf{Y}} = \mathbf{X}\hat{\boldsymbol{\beta}} + \hat{\mathbf{u}}$, and then used it to estimate the root of the mean squared error (RMSE) of each fold as $RMSE_{fold} = \frac{1}{m}\sum_{i=1}^{m}(y_i - \bar{y})^2$, and calculated the final RMSE as the average across all CV folds. For each data set, similarity matrix algorithm (LDAK/GCTA or GeSi), and data type (WGS, LD-pruned or RS), we found the $\alpha$ value that minimized the average RMSE. Similarly, we estimated the Pearson correlation coefficient between the true genetic value and the predicted genetic value (simulated data only), and between the true phenotype and predicted phenotype (simulated and BioMe data).
